# Mutations in *Bcl9* and *Pygo* genes cause congenital heart defects by tissue-specific perturbation of Wnt/β-catenin signaling

**DOI:** 10.1101/249680

**Authors:** Claudio Cantù, Anastasia Felker, Dario Zimmerli, Elena Chiavacci, Elena María Cabello, Lucia Kirchgeorg, Tomas Valenta, George Hausmann, Jorge Ripoll, Natalie Vilain, Michel Aguet, Konrad Basler, Christian Mosimann

**Author notes:** contributed equally to this work (Claudio Cantù, Anastasia Felker, Dario Zimmerli).

## Abstract

Genetic alterations in human *BCL9* genes have repeatedly been found in congenital heart disease (CHD) with as-of-yet unclear causality. BCL9 proteins and their Pygopus (Pygo) co-factors can participate in canonical Wnt signaling via binding to β-catenin. Nonetheless, their contributions to vertebrate heart development remain uncharted. Here, combining zebrafish and mouse genetics, we document tissue-specific functions in canonical Wnt signaling for BCL9 and Pygo proteins during heart development. In a CRISPR-Cas9-based genotype-phenotype association screen, we uncovered that zebrafish mutants for *bcl9* and *pygo* genes largely retain β-catenin activity, yet develop cardiac malformations. In mouse, both systemic and lineage-specific loss of the Pygo-BCL9-β-catenin complex caused heart defects with outflow tract malformations, aberrant cardiac septation and valve formation, and compact myocardium hypoplasia. Mechanistically, these phenotypes coincide with transcriptional deregulation during heart patterning, and Pygo2 associates with β-catenin at *cis*-regulatory regions of cardiac genes. Taken together, our results establish BCL9 and Pygo as tissue-specific β-catenin co-factors during vertebrate heart development. Our results further implicate alterations in *BCL9* and *BCL9L* in human CHDs as possibly causative.

## Introduction

Congenital heart diseases (CHD) are the most frequent human birth defects (affecting ca. 0.8% of live births) and can result in perinatal lethality or life-long complications despite increasingly successful surgical interventions^1^. Understanding the molecular details of heart development is imperative for discovering the causes of CHD. The heart is the first functional organ in the developing vertebrate embryo. It initially forms as a linear tube from the anterior lateral plate mesoderm (LPM) before undergoing extensive remodeling and looping^2^. Addition of second heart field (SHF)-assigned late-differentiating LPM expands the heart and triggers the formation of the right ventricle and atria in mammals^3^. Migrating cardiac neural crest cells (CNC) have further been implicated in contributing to heart formation, in particular by populating the outflow tract (OFT)^4^.

Mutations in various genomic loci, in particular in cardiac transcription factor genes including *GATA4*, *TBX5*, *MSX1*, or *HAND1/2*, have been linked to familial CHD^5^ and have become functionally associated with CHD phenotypes by genetic studies in model organisms^6^. Nonetheless, the predominant number of spontaneous CHD cases with identified sequence alterations do not feature clear recessive loss-of-function mutations, but are frequently heterozygous for mutations or harbor copy number variations (CNVs) with amplifications and deletions of genomic regions encompassing several genes^7,8^. Markedly, while heterozygous perturbations seem sufficient to cause CHDs in humans, mutations in CHD-implicated genes in mouse and zebrafish frequently act as recessive trait^9,10^. These observations indicate that human cardiac development might be particularly sensitive to alterations in gene dosage.

Canonical Wnt signaling contributes to numerous developmental processes, including heart formation, by controlling the nuclear role of β-catenin as transcriptional regulator and its cooperation with tissue-specific transcription factors^11,12^. β-catenin activity directs proliferation and cell fate commitment at different stages and in different cell types during heart formation. During gastrulation, canonical Wnt signaling promotes mesoderm formation, while its inhibition is subsequently required for LPM induction into early cardiac precursors^13^. Early cardiomyocytes display a robust proliferative response to canonical Wnt signaling and, conversely, accelerated differentiation upon pathway inhibition^14^. Several phases of canonical Wnt signaling are guiding the interplay of endocardial and myocardial layers to form functional heart valves^13,15^. Canonical Wnt signaling has also been linked to enhanced proliferation of SHF cells and to the control of CNC cells that both contribute to the mammalian OFT^16–19^. How canonical Wnt signaling elicits tissue-specific responses through the seemingly universal action of β-catenin in target gene control remains incompletely understood.

Nuclear β-catenin orchestrates canonical Wnt target-gene expression by recruiting a host of co-factors to Wnt-responsive elements (WREs) occupied by TCF/LEF transcription factors^20,21^. In *Drosophila*, the coupling of the histone code-reader plant homology domain (PHD) finger protein Pygopus (Pygo) via the adaptor protein Legless (Lgs) to the β-catenin N-terminal Armadillo repeats is necessary for virtually all canonical Wnt signaling-dependent responses^22–24^. Vertebrate genomes commonly harbor two paralogs of *lgs*, *Bcl9* and *Bcl9l* (*Bcl9/9l*), and two of *pygo*, *Pygo1* and *Pygo2* (*Pygo1/2*). Morpholino antisense experiments and overexpression have suggested functions in β-catenin-associated axis formation during gastrulation in Xenopus and zebrafish^25,26^. Nonetheless, genetic loss-of-function in the mouse has questioned their relevance for canonical Wnt signaling in mammals: neither the compound deletion of *Bcl9/9l* nor the deletion of *Pygo1/2* causes a general abrogation of canonical Wnt signaling in mouse development^27–30^. Mouse mutant analysis has further emphasized that BCL9 and Pygo proteins have evolved β-catenin-independent and even cytoplasmic functions in mammals, including the genetic interaction with Pax6 during eye lens formation and cytoplasmic functions in enamel formation during tooth development^29–31^. Biochemical evidence further suggests that Pygo-BCL9 act as connecting factors of chromatin-engaged complexes^23^. Whether and to what extent BCL9/9L and Pygo1/2 at all contribute to β-catenin-dependent transcription during mammalian development remains to be resolved^27–29,31^. Alterations in *BCL9* have been repeatedly associated with disease states; most notably, CNVs with locus gains and losses of the human chromosome *1q21.1* locus, to which *BCL9* maps to, are repeatedly found in cases of CHD^32–38^, while mutant *BCL9L* has been linked to heart-affecting heterotaxia^39^. If and what role BCL9 plays during heart formation remains unaddressed.

Here, we provide evidence that BCL9 and Pygo proteins contribute as tissue-specific mediators of β-catenin activity to vertebrate heart formation. Following a CRISPR-Cas9-based phenotype screen of CHD-implicated candidate genes in zebrafish, we uncovered that mutating the Pygo-BCL9-β-catenin complex causes impaired heart formation. Late developmental defects of *bcl9* and *pygo1/2* mutants are phenocopied by selective chemical inhibition of the BCL9-β-catenin interaction when perturbation is initiated before, during, or after gastrulation. In the mouse heart, we uncovered septation, OFT, and valve defects by both constitutive and heart-specific conditional loss of *Bcl9/9l* or *Pygo1/2*, or by simultaneous impairment of the BCL9/9L-β-catenin and BCL9/9L-PYGO2 interactions. Transcriptome and chromatin binding analyses established that the PYGO-BCL9-β-catenin complex contributes to the regulation of key factors involved in cardiac patterning and OFT and heart valve formation, including Pitx2, Hand2, Prrx1 and Msx1. Collectively, our results provide functional and molecular evidence for a conserved tissue-specific function of the BCL9 and Pygo factors in canonical Wnt signaling in vertebrates. These findings further suggest a causative link between human CHDs and copy number changes in *BCL9* at *1q21.1* and mutations reported in *BCL9L* that can potentially perturb the expression of heart-specific canonical Wnt target genes during cardiac development.

## Results

### BCL9 and Pygo perturbations cause developmental heart defects in zebrafish

To identify regulators of vertebrate heart formation implicated in chromosomal amplifications and deletions found in human CHDs, we performed an injection-based candidate gene screen in zebrafish using maximized CRISPR-Cas9-mediated mutagenesis in F0 crispants^40^ (Figure 1a). We targeted candidate genes with individual sgRNAs by injection of Cas9 ribonucleoprotein complexes into one-cell stage zebrafish embryos, and screened the resulting crispants for impaired heart development (Figure 1a). These efforts revealed that mutations in *bcl9* lead to pericardial edema within the first 120 hpf of zebrafish development (Figure 1a). We established heterozygous mutant zebrafish strains for both BCL9 family genes *bcl9* and *bcl9l*, featuring frameshift deletions in-between the coding sequences for HD1 and HD2, the domains of BCL9 proteins that convey functional interaction with the interactors Pygo and β-catenin, respectively^22,29^. We refer to these new alleles as *bcl9^Δ29^* and *bcl9l^Δ4^*: if not null, their BCL9 protein products have impaired association with β-catenin (Figure 1b, Supplementary Figure 1). Heterozygous *bcl9^Δ29^* and homozygous *bcl9l^Δ4^* zebrafish were viable and fertile with no obvious phenotypes (observed for more than 4 generations) (Supplementary Figure 1). Contrary to previous conclusions based on morpholino-mediated knockdown^25^, this observation suggests that BCL9L does not contribute any essential functions in zebrafish under laboratory conditions.

**Figure 1:**
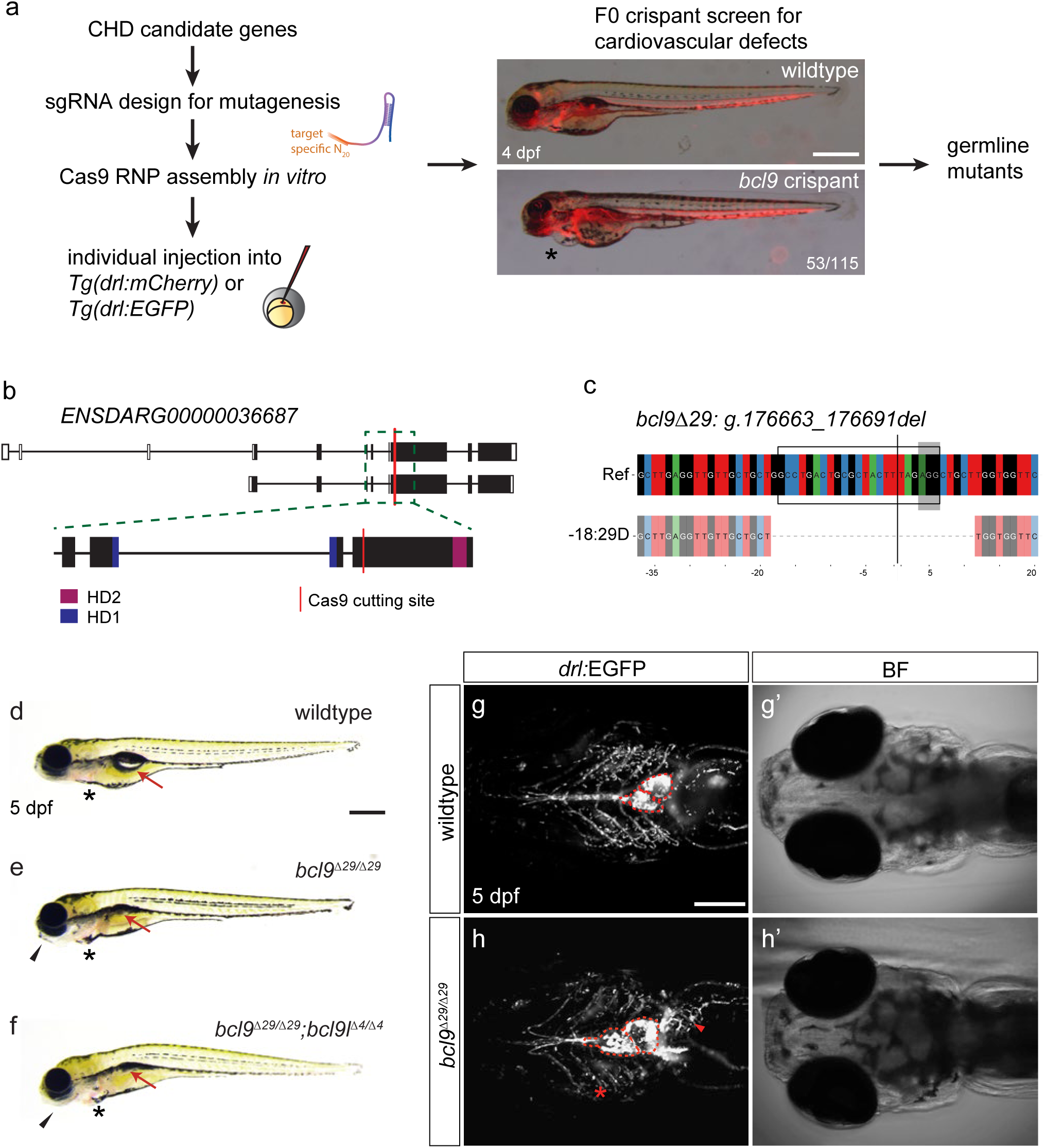
Mutations in *bcl9* lead to severe cardiac and craniofacial defects in zebrafish. (**a**) A CRISPR-Cas9-mediated mutagenesis screen for cardiac defects (asterisk) identified *bcl9* as a potential new factor involved in heart morphogenesis. *bcl9* crispants have heart looping defects as visible in *drl*:mCherry transgenics leading to cardiac edema (aterisks), lateral view, anterior to the left. (**b**) Schematic representation of the *bcl9* gene locus. The sgRNA was designed to target the coding exon 6 between HD1 and HD2 of the zebrafish *bcl9* gene. The locus is represented as per genome annotation Zv10 with two isoforms that differ in the first coding exon and the untranslated regions (UTRs). The green dotted box represents a zoomed region of the gene locus, with the red line representing the location of the sgRNA used for mutagenesis; black boxes mark coding exons (CDS), white boxes mark UTRs, the blue boxes represent the part of the CDS that will contribute to HD1 and purple boxes to HD2. Note that the schematic is not to scale. (**c**) CrispRVariants panel plot depiction of the germline allele with a 29 bp deletion at the Cas9 cutting site. Top shows genomic reference, while the *bcl9:g.176663_176691* (*bcl9^Δ29^*) allele is shown below. *bcl9^Δ29^* results in an out-of-frame deletion introducing a frameshift in the CDS. The black box indicates the exact position of the sgRNA sequence, the grey shaded box the 5’-NGG-3’ PAM sequence, and the black line the predicted Cas9-induced double-strand break position. (**d-f**) Brightfield images of 5 dpf homozygous *bcl9^Δ29^* and *bcl9^Δ29^*;*bcl9l^Δ4^* larvae and their wildtype siblings, lateral views, anterior to the left. Mutant larvae show heart looping defects that result in cardiac edema (asterisks). Moreover, mutant larvae do not inflate their swim bladder (arrows) presumably due to a failure in gasping air because of craniofacial malformations (black arrow heads). (**g-h**) SPIM images of *drl*:EGFP-expressing wildtype and homozygous *bcl9^Δ29^* larvae, ventral views, anterior to the left. Mutant larvae show craniofacial vasculature (asterisks and arrow head) and heart looping (dotted line outlines the atrium and ventricle) defects. Scale bars, 200 µm (**g-h**), 500 µm (**a,d-f**).

Zygotic-mutant embryos homozygous for *bcl9^Δ29^* developed seemingly normal until 48 hpf and did not show any misregulation of early cardiac markers at 24 hpf (Supplementary Figure 2). Between 56-72 hpf, while the hearts of homozygous *bcl9^Δ29^* mutants are beating and pattern into an atrial and ventricular chamber, we observed a variable incidence and expressivity of pericardial edema (38% ± 12) and misexpression of the cardiac valve marker *vcana* in mutant embryos (Supplementary Figure 3). By 5 days post fertilization (dpf), homozygous *bcl9^Δ29^* larvae showed highly penetrant craniofacial and cardiac defects (n=181/879, 20.6% compared to Mendelian 25%, N=6) (Figure 1d,e): lightsheet imaging of homozygous *bcl9^Δ29^* mutants revealed an underdeveloped cardiopharyngeal vasculature, vascular defects at the cardiac inflow tract, a smaller OFT, and perturbed heart looping with misaligned atrium and ventricle when compared to wildtype or heterozygous siblings (Figure 1g,h). Alcian blue staining to visualize cartilaginous craniofacial elements in zygotic *bcl9^Δ29^* mutants further documented deformed pharyngeal skeletons with abnormal Meckel’s and palatoquadrate cartilage development and fusion defects of the ceratohyal and ceratobranchial 1 cartilage, with high penetrance (Supplementary Figure 4). Homozygous *bcl9^Δ29^*–mutant larvae also failed to properly inflate their swim bladder and died at 11-12 dpf. Homozygous *bcl9^Δ29^*;*bcl9^Δ4^* double-mutant zebrafish embryos looked indistinguishable from homozygous *bcl9^Δ29^* mutants (Figure 1d–f), confirming the non-essential function of *bcl9l,* and suggesting no significant genetic compensation^41^ by the paralog *bcl9l* in *bcl9^Δ29^* mutants.

BCL9/9L proteins act during *Drosophila* and vertebrate development via functional coupling with the histone-code reader Pygo^22,42–44^. To test for a potential role of Pygo1/2 during zebrafish heart development, we generated *pygo1* and *pygo2* loss-of-function mutant alleles. These alleles harbor frameshift mutations within the essential NH2-terminal homology domain (NHD) and result in a premature Stop codon before the BCL9-binding C-terminal PHD domain; we refer to these alleles as *pygo1^Δ5^* and *pygo2^Δ1^* (Figure 2a–c). Homozygous *pygo1^Δ5^* and homozygous *pygo2^Δ1^* zebrafish embryos were indistinguishable from wildtype siblings throughout development, and homozygous *pygo1^Δ5^* and homozygous *pygo2^Δ1^* zebrafish develop to fertile adults (Supplementary Figure 5). In contrast, embryos homozygous for *pygo2^Δ1^* and compound hetero– or homozygosity for *pygo1^Δ5^* displayed cardiac and craniofacial cartilage defects that recapitulated those observed in homozygous *bcl9^Δ29^* zebrafish mutants (Supplementary Figures 4 and 5). Taken together, these data reveal select post-gastrulation defects and sensitivity of cardiac development to functional BCL9 and Pygo levels in zebrafish.

**Figure 2:**
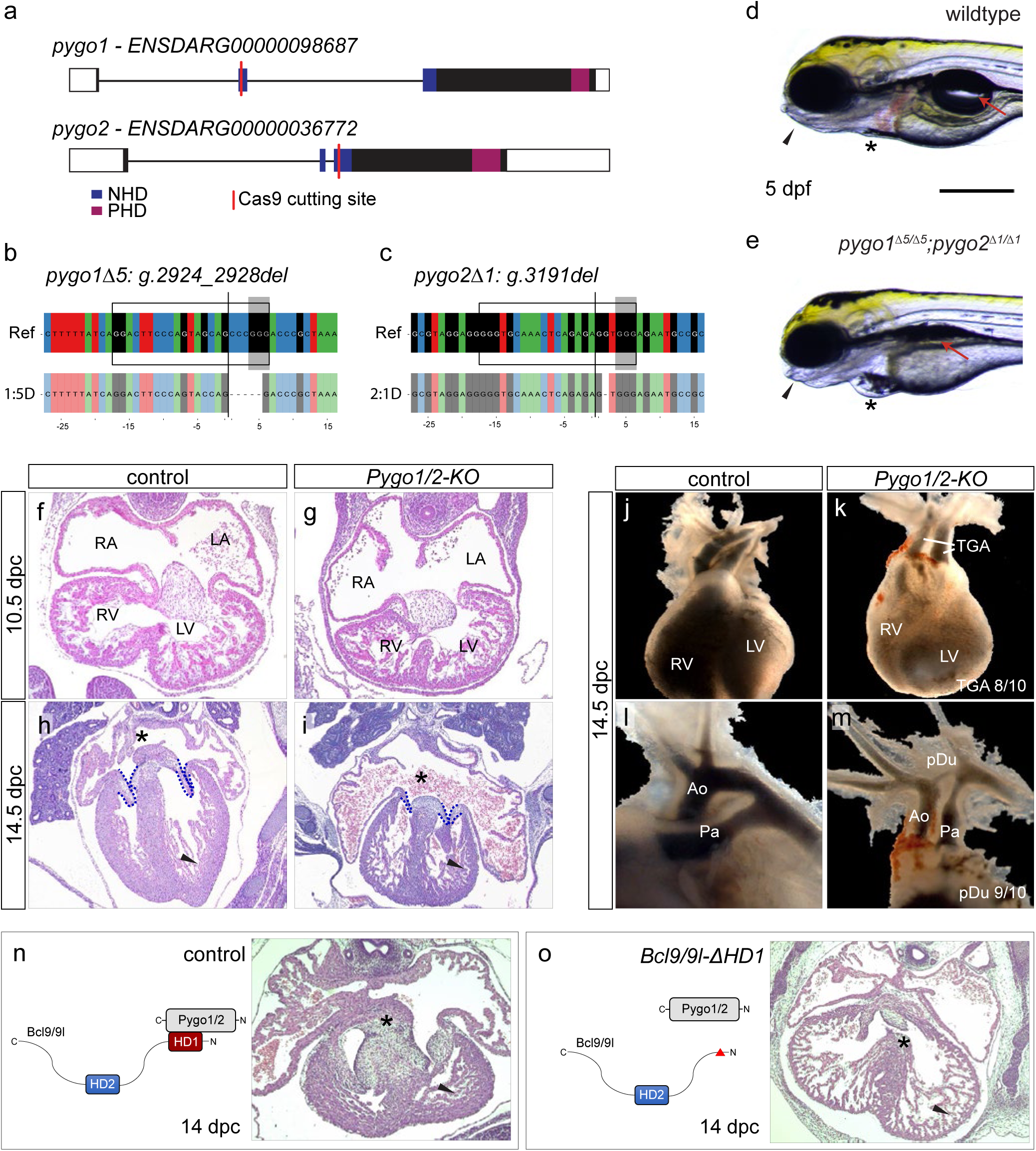
*Pygo1/2* mutant zebrafish and mouse embryos develop cardiac malformations reminiscent of CHDs. (**a**) Schematic representation of the zebrafish *pygo1* and *pygo2* genes with annotated NHD and PHD domains and the Cas9 cutting site to generate mutants. Gene locus represented as per genome annotation Zv10 with the main isoforms of both genes shown. See description of (**1b**). (**b-c**) CrispRVariants panel plot depictions of the germline alleles with a 5 bp or 1 bp deletion in *pygo1* and *pygo2*, respectively. Top shows genomic reference with the *pygo1Δ5: g.2924_2928del* and *pygo2Δ1: g.3191del* alleles shown below. Both alleles result in an out-of-frame deletion introducing a frameshift in the CDS. See description of (**1c**). (**d-e**) Brightfield images of live *pygo1^Δ5^*;*pygo2^Δ1^* double mutants reveal cardiac edema (asterisks) and craniofacial defects (arrow heads), and aberrant swim bladder inflation (arrows) as detected in *bcl9^Δ29^* mutants, lateral views, anterior to the left. (**f-i**) Haematoxylin/eosin stained sagittal sections of the murine heart at 10.5 (**f,g**) and 14.5 dpc (**h,i**). At 10.5 dpc the development of the heart and heart cushion was still largely normal in the mutants. At 14.5 dpc the mutants displayed markedly smaller and thinner valves (dashed outline) and compact ventricular myocardium (arrow head), highly dilated atria, and the atrial septum was missing (asterisks). RA, right atrium; LA, left atrium; RV, right ventricle; LV, left ventricle. (**j-m**) Gross anatomical view of heart and great vessels at 13.5/14.5 dpc as revealed by India ink injection. While normally the aorta (Ao) arises from the left (LV) and the pulmonary artery (Pa) from the right ventricle (RV), mutants had a classic transposition of the great arteries (TGA) and a patent ductus arteriosus (pDu). (**n-o**) Schematic representation of the Bcl9/9l-Pygo1/2 interaction (n) and the molecular configuration of this interaction when the HD1 domain in Bcl9/9l is deleted in *Bcl9/9l-ΔHD1* mice (**o**); and heart sections at 14 dpc, stained with Haematoxylin/eosin. The abrogation of this interaction leads to a delayed chamber septation (asterisks), hypoplastic myocardium (arrow head), and valve deficiency. Scale bar, 250 µm (**d,e**).

### BCL9 and PYGO are required for proper heart development in the mouse

In mouse, compound loss-of-function of *Pygo1* and *Pygo2* (*Pygo1/2*) leads to embryonic lethality at 13.5/14.5 days post-coitum (dpc)^27,28^. Analysis of these mutants revealed that *Pygo1/2*-mutant mouse embryos develop severe heart defects between 10.5 and 14.5 dpc: histology and 3D reconstruction of the heart documented shorter and thinner atrio-ventricular valve leaflets, dilated atria, hypoplastic ventricular myocardium, and aberrant chamber septation (Figure 2f–i, Supplementary Videos 1 and 2, Supplementary Figure 6). In addition, *Pygo1/2*-mutant embryos displayed prominent OFT anomalies such as transposition of the great arteries (TGA, penetrance of 80%, n=10; Figure 2j,k) and a hypotrophic ductus arteriosus (penetrance of 90%, n=10; Figure 2l,m).

We subsequently tested the requirement of PYGO1/2 binding to BCL9/9L in heart formation by making use of *Bcl9/9l* alleles carrying a deletion in the region encoding the PYGO1/2-binding domain HD1 (*Bcl9/9l-ΔHD1*)^29^. *Bcl9/9l-ΔHD1-*homozygous mouse embryos died at the same stage as *Pygo1/2* mutants (13.5-14.5 dpc), and histological sections revealed heart defects resembling those observed upon *Pygo1/2* loss (Figure 2n,o). This result indicates that the cardiac defects observed in our mutant mice arise from perturbing the cooperative action of BCL9/9L and PYGO1/2.

Taken together, our analyses in mouse and zebrafish establish that perturbation of BCL9 and Pygo function cause defective cardiac development in two evolutionarily distant vertebrates.

### The BCL9-Pygo complex drives Wnt/β-catenin signaling in the developing heart

Due to the β-catenin-independent roles of BCL9 and Pygo proteins^28–31^, we cannot formally link the phenotypes resulting from our mutants to defective canonical Wnt signaling. Since BCL9/9L can act as linker proteins connecting β-catenin and PYGO^45^, we aimed at specifically testing the concurrent requirement of connecting these interaction partners.

We generated mice trans-heterozygous for alleles of *Bcl9/9l* in which either the HD1 (PYGO binding) or the HD2 (β-catenin binding) domains are deleted (Figure 3a,b)^29^. Double-heterozygous mice carrying a deletion in HD1 (*Bcl9^ΔHD1/+^*; *Bcl9l^ΔHD1/+^*, referred to as *Bcl9/9l-ΔHD1/+*) or in HD2 (*Bcl9^ΔHD2/+^*; *Bcl9l^ΔHD2/+^*, referred to as *Bcl9/9l-ΔHD2/+*) are viable and fertile. Crosses between *Bcl9/9l-ΔHD1/+* and *Bcl9/9l-ΔHD2/+* must lead to a trans-heterozygous configuration (in 1/16 embryos) in which both domain deletions are present (*Bcl9^ΔHD1/ΔHD2^*; *Bcl9l^ΔHD1/ΔHD2^*, referred to as *Bcl9/9l-ΔHD1/ΔHD2*). The resulting protein products can form either BCL9-β-catenin or BCL9-PYGO complexes, but not the full tripartite transcriptional complex (Figure 3b), allowing us to test the developmental requirement of the PYGO-BCL9-β-catenin complex.

**Figure 3:**
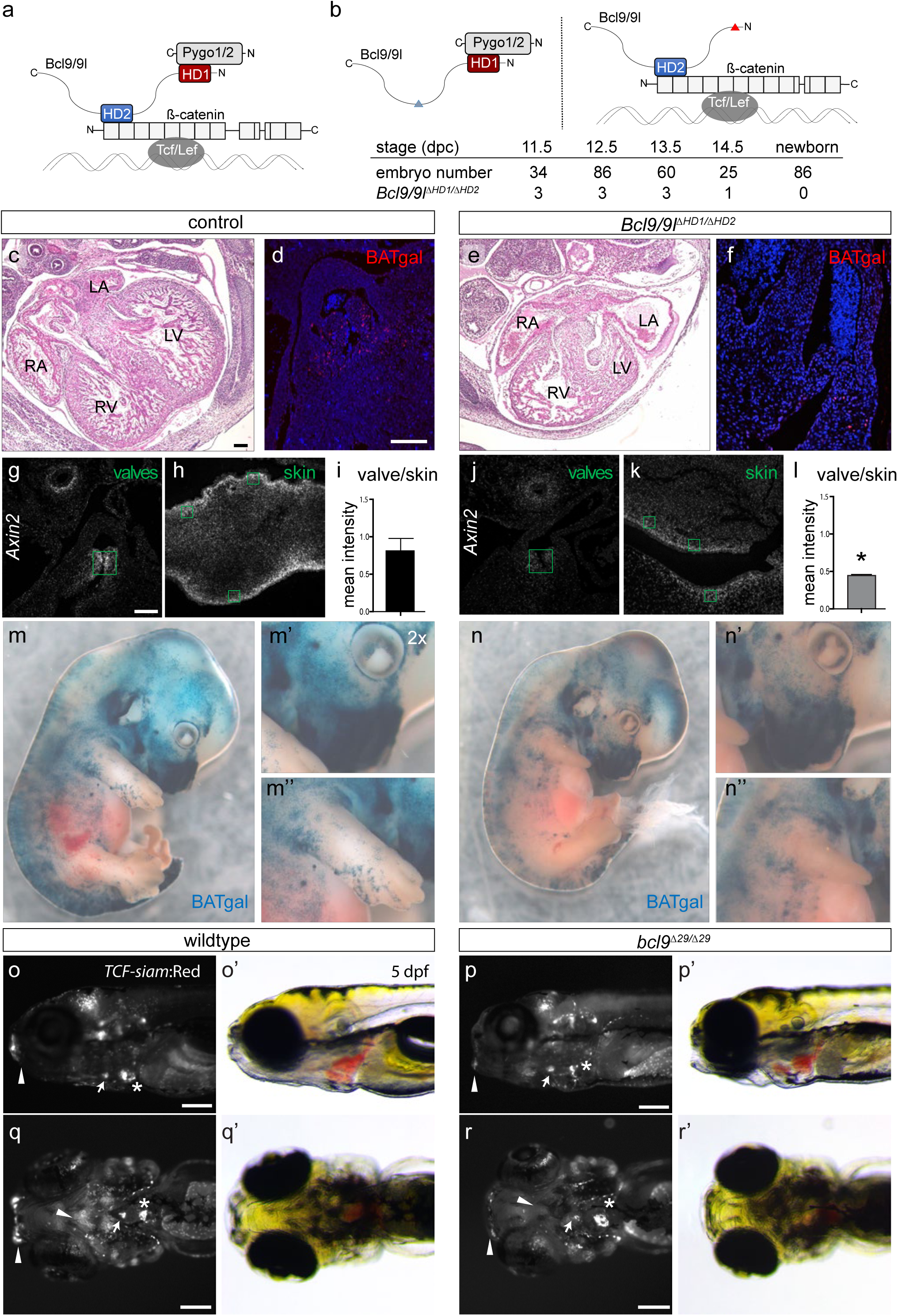
The BCL9-Pygo complex drives Wnt/β-catenin transcription during heart development. (**a**) Schematic representation of the β-catenin transcriptional complex. Bcl9/9l simultaneously interacts with Pygo1/2 and β-catenin via the domains HD1 and HD2, respectively. (**b**) In *Bcl9/9l-Δ1/Δ2* mice, no Bcl9/9l molecule can simultaneously bind Pygo1/2 and β-catenin; the deletion of the HD2 or HD1 domains abrogates Bcl9/9l binding to β-catenin (left panel) or Pygo1/2 (right panel), respectively. *Bcl9/9l-Δ1/Δ2* embryos are never found after embryonic stage 14.5 dpc (table). (**c-f**) Haematoxylin/eosin staining on heart sections of control (**c**) and *Bcl9/9l-Δ1/Δ2* embryos (**e**) at 13.5 dpc. *Bcl9/9l-Δ1/Δ2* embryos have pronounced heart defects, with thinned myocardium of the ventricular walls, malformations of the forming septum, the AV valves and OFT. The expression of BATgal in vivo Wnt reporter in the OFT region is markedly reduced in *Bcl9/9l-Δ1/Δ2* mutant (**f**) compared to control (**d**) hearts. RA, right atrium; LA, left atrium; RV, right ventricle; LV, left ventricle. (**g-l**) *Axin2* expression in wildtype and *Bcl9/9l-Δ1/Δ2* OFT valves (**g,j**) and skin at 13.5 dpc. In g and j, green boxes are drawn around transversal sections of individual valves of the OFT. (**h,k**) with quantification of fluorescence intensity levels (**i,l**). *Axin2* expression is strongly reduced in the valves of *Bcl9/9l-Δ1/Δ2* embryos (**j**), while reduction in the skin is mild (**k**). The quantification of the ratio between *Axin2* expression in the valves and the skin in the different genotypes reveals a significant reduction of *Axin2* expression in Bcl9/9l-Δ1/Δ2 valves (**i,l**). In h and k, green boxes mark the regions of skin covering the embryonic limbs, from which signal intensity was quantified. (m-n) In vivo BATgal reporter expression in 13.5 dpc wildtype (**m**) and *Bcl9/9l-Δ1/Δ2* mutant (n) embryos. Mutant embryos have a slightly decreased reporter expression, but BATgal expression is generally retained in most tissues. Reduced *BATgal* activity is observed in the craniofacial region (**m’,n’**) and in the forelimbs (**m”,n”**). Loss of canonical Wnt signalling transcription in forelimbs is accompanied by severe developmental limb defects in *Bcl9/9l-Δ1/Δ2* embryos (compare **m”** with **n”**). (**o-r**) Fluorescent and brightfield images of TCF-siam:Red *bcl9^Δ29^* and wildtype zebrafish larvae, lateral (**m,n**) and ventral (**o,p**) views, anterior to the left. *TCF* reporter activity is specifically reduced in the cardiac OFT (arrow) and craniofacial apparatus (arrow heads), and altered in the atrio-ventricular valve (asterisks). Scale bars, 100 µm (**c-l**), 200 µm (**o-r**).

From these crosses, we never recovered *Bcl9/9l-ΔHD1/ΔHD2* pups (Figure 3b, bottom table), indicating embryonic lethality. Even though at lower numbers than the expected Mendelian ratio, *Bcl9/9l-ΔHD1/ΔHD2* embryos did reach the 13.5-14.5 dpc stage (Figure 3b) and displayed heart defects including ventricular myocardium hypoplasia and smaller and thinner valve leaflets, recapitulating the defects observed upon *Pygo1/2* loss (Figure 3c-f). These findings indicate that BCL9/9L is required to physically connect to both PYGO1/2 and β-catenin for proper heart development.

Further relating these defects to perturbed canonical Wnt signaling was the reduced expression of the *in vivo BATgal* reporter (sensing nuclear β-catenin activity) in *Bcl9/9l-ΔHD1/ΔHD2* mutants in cardiac valve progenitors, a region with active canonical Wnt signaling^46^ (Figure 3d,f). Single molecule mRNA *in situ* hybridization of *Axin2*, a prototypical pan-canonical Wnt target gene, confirmed that decreased Wnt signaling occurred in compound mutants in valve progenitors but not in other tissues such as the skin at time of analysis^47^ (Figure 3g-l). In addition to cardiac defects, we observed severely underdeveloped limbs (Figure 3m–n), and skeletal malformations (Supplementary Figure 7) including shortened radius and ulna bones, incorrect specification of digit number (Supplementary Figure 7b,e), and bifid ribs (Supplementary Figure 7c,f). Notably, while broadly unaffected in *Bcl9/9l-ΔHD1/ΔHD2* embryos, the reduction of *BATgal* expression occurred, in addition to the cardiac valve progenitors, mainly in the other malformed tissues including craniofacial structures (Figure 3m',n') and the developing forelimbs (Figure 3m”,n”). In *bcl9^Δ29^*-mutant zebrafish at 5 dpf, the pattern and strength of the canonical Wnt signaling reporter *Tg(7xTCF-Xla.Siam: nlsmCherry)*^*ia5*^ (referred to as *TCF-siam:Red*^48^) were affected in the atrio-ventricular valve, in the OFT, and in craniofacial structures (Figure 3o–r). These data emphasize that disconnecting β-catenin from the Pygo-BCL9 module during both mouse and zebrafish development does not cause systemic perturbation of canonical Wnt signaling, yet has selective impact on isolated cell types and processes.

To independently perturb the protein-protein interaction between BCL9 and β-catenin in zebrafish, we used the chemical compound LH-2-40 that acts as selective inhibitor of the BCL9-HD2 interaction with the β-catenin Arm repeats 1-2^49^. We treated wildtype zebrafish embryos with LH-2-40 at concentrations ranging from 1 to 50 µM at distinct developmental time-points: 4 cell-stage, shield stage, and 18 somite stage, respectively. Pre-gastrulation– and gastrulation-stage treatment with LH-2-40 did not induce any observable gastrulation defects (Figure 4a,b); in contrast, at 5 dpf, embryos treated with LH-2-40 at any prior time point recapitulated the *bcl9^Δ29^*-mutant phenotypes in a dose-dependent manner (Figure 4c–h). Consistent with our genetic observations, LH-2-40-treated embryos displayed altered *TCF-siam:Red* expression in the atrio-ventricular valve and in craniofacial structures at 3 and 5 dpf (Figure 4i–l). In addition, when treating embryos obtained from *bcl9^Δ29^* heterozygous incrosses, we observed a higher penetrance of strong phenotypes resembling those observed in *bcl9^Δ29^* homozygous larvae (Supplementary Figure 8a–f). Akin to wildtype treatments (Figure 4a,b), however, we never observed gastrulation defects. Notably, in addition to the craniofacial and cardiac defects found in *bcl9^Δ29^* mutants, LH-2-40 treatment frequently caused pectoral fin defects, characterized by angled fin position and deformations of the interskeletal disc (Figure 4l, Supplementary Figure 8g,h).

**Figure 4:**
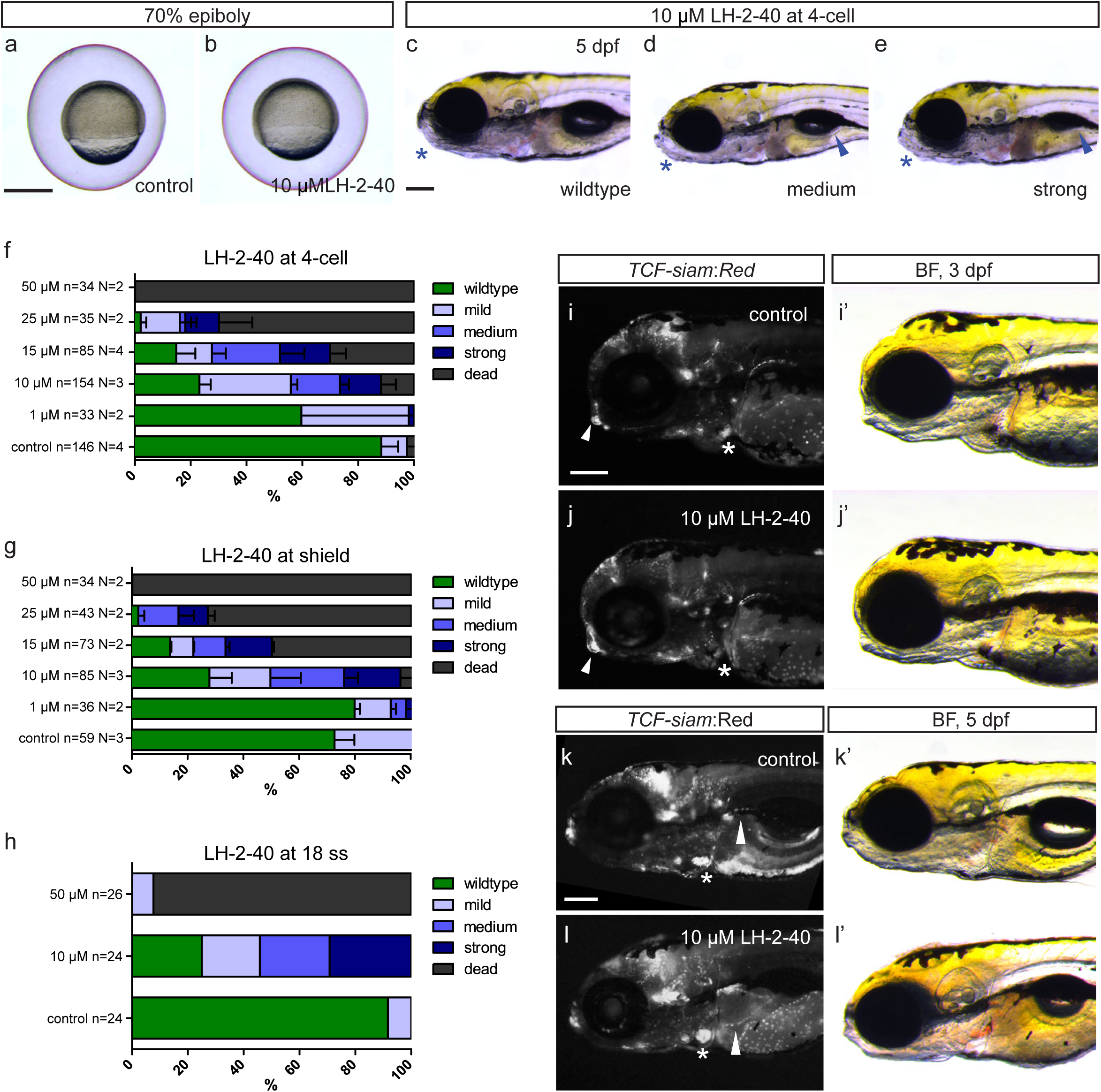
Chemical manipulation of zebrafish heart development by Bcl9-β-catenin inhibitor LH-2-40. (**a-b**) Treatment with 10 µM LH-2-40 from 2-4 cell stage does not result in gastrulation defects, lateral views. (**c-e**) Brightfield images of 5 dpf old DMSO controls (**c**) and representative larvae treated with 10 µM LH-2-40 from 4-cell stage on (**d-e**) as assessed for phenotype classes quantified in (**f-h**), lateral views, anterior to the left. Phenotypes in LH-2-40 treated embryos become visible at 5 dpf, comparable to phenotypes observed in *bcl9^Δ29^* mutants. LH-2-40 treated embryos show a variable phenotype expressivity with mild to strong swim bladder inflation defects (arrow heads **d,e**) caused by variable craniofacial defects (asterisks d,e, compare to control in **c**). (**h-j**) Similar dose-dependent phenotype penetrance and expressivity can be observed after Bcl9 inhibition at 4-cells, shield and 18 ss stage suggesting that the observed phenotypes result from Bcl9 function in craniofacial and heart development after somitogenesis. Mild phenotypes are characterized by mild craniofacial and swim bladder inflation defects. Both phenotypes are more pronounced in the medium phenotype class (see **d**). Strong phenotypes are characterized by strong craniofacial defects, a complete failure to inflate the swim bladder (see **e**). (**i-l**) Fluorescent and brightfield images of *TCF:siamRed* Bcl9 inhibited (**j,l**) and DMSO treated (**i,k**) wildtype siblings at 3 and 5 dpf, lateral views, anterior to the left. *TCF* reporter activity is severely reduced in the atrio-ventricular valve at 3 dpf (asterisks i,j) and altered in the craniofacial cartilage (arrow heads **i,j**). At 5 dpf, TCF is miss-expressed in the atrio-ventricular valve (asterisks **k,l**) and fins (arrow heads **k,l**). Scale bars, 200 µm (**c-e**,**i-l**), 500 µm (**a,b**).

While we cannot rule out influences by Wnt/β-catenin-independent functions of Bcl9 and Pygo, our results are consistent with their joint requirement for β-catenin-dependent signaling during selective vertebrate developmental processes, including heart formation.

### Mesodermal and neural crest-derived cardiac progenitors are perturbed upon mutations in *BCL9* and *Pygo*

β-catenin-dependent target gene control is required at different stages of vertebrate heart development, including the initiation of the cardiac program within the LPM, chamber formation, and valve development^50^. We therefore aimed at capturing the gene expression changes that accompanied the embryonic defects caused by mutations in *Bcl9* and *Pygo*. Transcriptome analysis of anterior embryo structures affected by BCL9 perturbation (including heart, pharyngeal arches, pectoral fins, and craniofacial structures, Figure 5a) from *bcl9^Δ29^*-mutant zebrafish at 54 hours post fertilization (hpf) compared to wildtype siblings detected selective deregulation of 157 genes (83 genes down, FC <0.5; 74 genes up, FC>2.5) (Figure 5b; Supplementary Figure 9). The deregulated genes fall within different annotated gene expression categories, including heart, pharyngeal arches, and cranial neural crest (Supplementary Table 2). Moreover, several of the deregulated genes have been linked, in particular, to epithelial-to-mesenchymal transition (EMT) and to morphogenetic processes occurring during cardiac valve formation (Figure 5b, Supplementary Table 2). STRING analysis of our candidate gene list revealed medium-to-high confidence protein-protein interaction clusters related to cardiac valve formation (Figure 5c). While zebrafish do not form all structures found in the mammalian heart, this transcriptome profiling raised the possibility that BCL9, and by inference its connection with β-catenin and Pygo, are selectively required during specific aspects of vertebrate cardiac development. We therefore sought to complement our phenotype analysis of BCL9 and PYGO loss in cardiac development using conditional loss-of-function alleles in the mouse.

**Figure 5:**
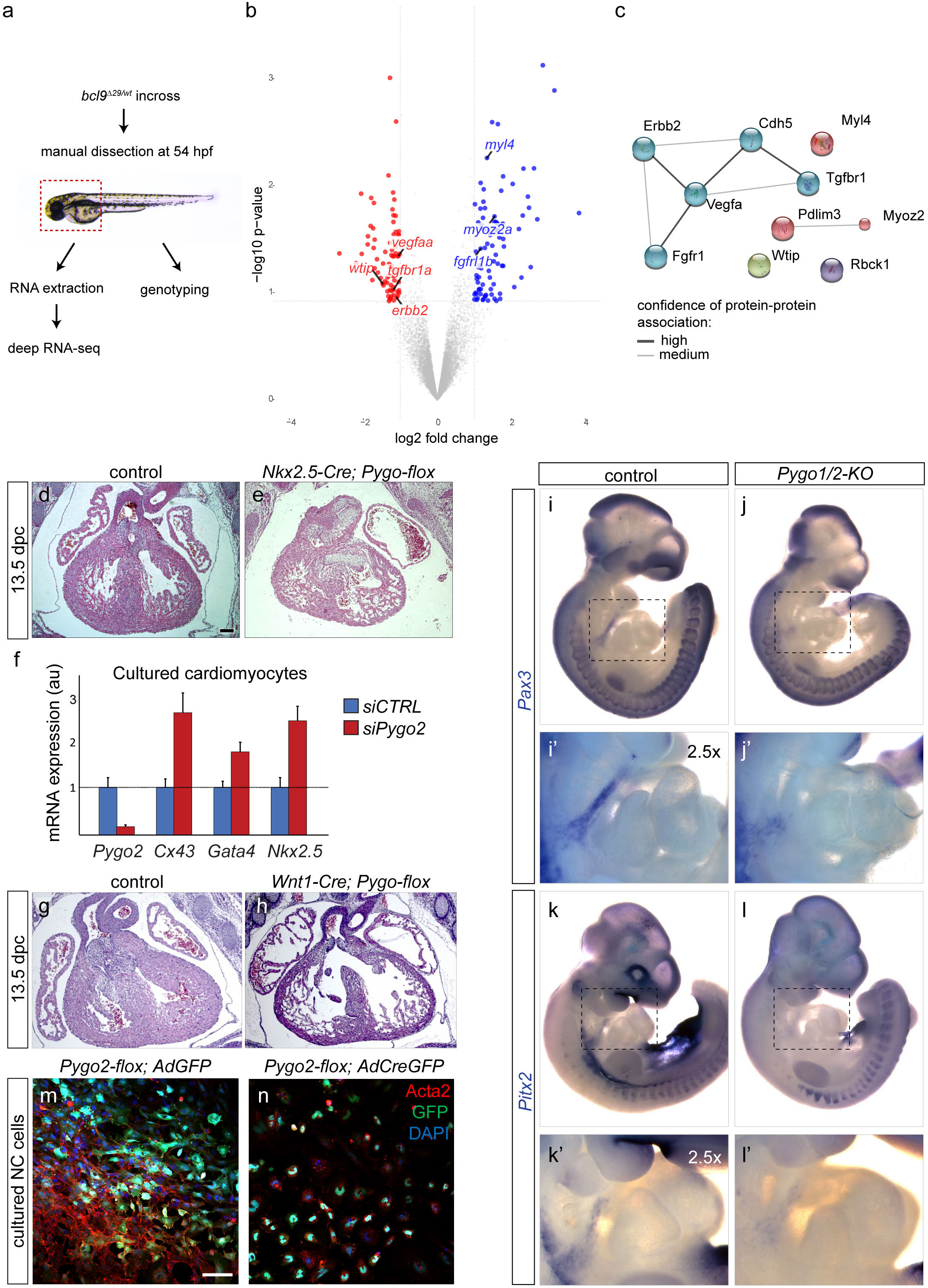
The Bcl9-Pygo complex acts in neural crest and cardiac heart progenitor cells. (**a**) Schematic representation of the RNA-seq approach used for *bcl9^Δ29^* mutant zebrafish embryos. RNA extraction was performed on manually dissected anterior structures of 54 hpf embryos as indicated by the dashed box. (**b**) Comparing three independent biological replicates revealed 157 differentially expressed (74 up– and 83 downregulated) genes between wildtypes and *bcl9^Δ29^* mutants as represented in the volcano plot. Genes associated with cardiac development are highlighted. (**c**) STRING analysis of mouse orthologues of de-regulated genes in *bcl9^Δ29^* zebrafish mutants with K-means clustering analysis (numbers of clusters 4). Five genes assemble in one cluster with medium to high confidence protein-protein interactions reflecting their interactive involvement in cardiac valve formation as reviewed from the literature. A second cluster reflecting of genes implicated in cardiomyocytes actin-myosin and sarcomere assembly assembles around upregulated genes. (**d,e**) Haematoxylin/eosin staining of heart section of control (**d**) and *Nkx2.5-Cre; Pygo1/2-flox* mouse embryos (**e**) at 13.5 dpc. (f) RT-qPCR on cultured embryonic cardiomyocytes indicates increased expression of cardiac differentiation markers when *Pygo2* is downregulated via siRNA. (**g,h**) Haematoxylin/eosin staining of heart section of control (**h**) and *Wnt1– Cre*; *Pygo1/2-flox* embryos (**i**) at 13.5 dpc. (**i-l**) Whole mount ISH showing expression of Pax3 (**i,j**) and Pitx2 (**k,l**) in representative control (**i,k**) and *Pygo1/2-KO* mutant (**j,l**) embryos at 10.5 dpc. Expression of the neural crest marker Pax3 is reduced in branchial arches (**i’,j’**) upon loss of *Pygo1/2*. *Pitx2* expression is drastically reduced in mutant embryos, particularly in periocular mesoderm, oral ectoderm, anterior somites (**k,l**), and branchial arches (**k’-l’**). (**m,n**) Cultured neural crest (NC) cells from *Pygo2-flox* mice were treated with Adeno viral particle expressing GFP as control (*AdGFP*, **m**) or Cre (*AdCreGFP*, **n**). Cre-mediated *Pygo2* recombination impaired in vitro NC differentiation in smooth muscle actin (Acta2, red) expressing cells. Scale bars, 100 µm.

To test for the contribution of mouse BCL9 and PYGO in LPM-derived heart progenitors, we combined the *Pygo-flox and Bcl9/9l-flox* strains with *Nkx2.5-Cre*, in which Cre recombinase is active in the expression domain of the cardiac homeobox gene *Nkx2.5* at early cardiac crescent stages (ca. 7.5 dpc)^51^. *Nkx2.5-*Cre-mediated *Pygo1/2* or *Bcl9/9l* recombination led to embryonic lethality around 13.5 dpc, accompanied by cardiac malformations including thinner myocardium and aberrantly enlarged atrio-ventricular valves (Figure 5d,e). The effect on the compact myocardium upon *Pygo1/2* or *Bcl9/9l* mutations is consistent with the role of canonical Wnt signaling in driving cardiomyocytes proliferation whilst preventing terminal maturation^52^. We confirmed that Wnt/β-catenin signaling, in this context, is dependent on PYGO proteins by performing a siRNA-mediated downregulation of *Pygo2* in cultured murine embryonic cardiomyocytes (Figure 5f): *Pygo2* knockdown in this system led to increased expression of the cardiomyocyte-specific genes *Gja1* (*Connexin43*), *Gata4*, and *Nkx2.5*, indicating enhanced differentiation^52^. Taken together, these experiments reveal that the BCL9-PYGO complex is required in mesodermal cardiac lineages, and that cardiac mesoderm-specific loss recapitulates aspects of the heart phenotypes induced by constitutive *Bcl9* and *Pygo* gene mutations.

In mammals and chick, CNC contributes to OFT development and septation^53,54^. We tested the requirement of BCL9 and PYGO in the cardiac CNC by combining the *Pygo-flox* and *Bcl9-flox* strains with the NC-specific *Wnt1-Cre* driver. *Pygo1^flox/flox^*; *Pygo2^flox/flox^*;*;Wnt1-Cre^Tg/*^* (referred to as *Wnt1-Cre;Pygo-flox*) and *Bcl9^flox/flox^*;*Bcl9l^flox/flox^*;*;Wnt1-Cre^Tg/*^* (referred to as *Wnt1-Cre;Bcl9/9l-flox*) displayed embryonic lethality at 13.5/14.5 dpc, and featured heart malformations that morphologically recapitulated the constitutive loss of *Pygo1/2* (Figure 5g,h). These results indicate that BCL9 and PYGO loss in CNCs is sufficient to perturb OFT valves and cardiac septation. The transcription factors Pax3 and Pitx2 are required for the migration and expansion of CNC cells at 10.5 dpc^16,55^; consistently, we observed decreased abundance of *Pax3* and *Pitx2* transcripts in migratory CNC cells in *Pygo1/2*-mutant embryos at 10.5 dpc, (Figure 5i-l). In addition, cultured primary neural crest cells isolated from the pharyngeal arches at 10.5 dpc from *Pygo-flox* mouse embryos, displayed signs of impaired non-neuronal differentiation when *Pygo1/2* conditional alleles were recombined via transduction of viral Adeno-CreEGFP particles. *Pygo2* downregulation led to impaired smooth muscle actin (SMA) fiber formation^56^ (Figure 5m,n, N=3), and reduced expression of the neural crest markers *Sox9* and *Osterix* as well as *Msx1,* which marks neural crest as well as mesodermal cardiac lineages of the OFT, right ventricle, and during valve formation^57,58^ (Supplementary Figure 10).

Taken together, these results establish that the BCL9-PYGO complex connected to β-catenin is required both in early mesodermal *Nkx2.5*-expressing cardiac progenitors and migrating CNC cells, the two key lineages whose interplay is crucial during heart development^59^.

### The tripartite Pygo-BCL9-β-catenin complex controls cardiac regulators

Our combined data provides evidence that the Pygo-BCL9-β-catenin complex contributes to canonical Wnt signaling activity during selected developmental processes in the vertebrate embryo, in particular in the heart. We next sought to gain insight into the target genes controlled by the PYGO-BCL9-β-catenin module in the CNC– and mesoderm-derived cardiac lineages. We dissected mouse embryonic branchial arches 3-6, through which CNC progenitor cells migrate dorso-ventrally toward the heart cushion and OFT structures^13^, together with the developing heart tube, from constitutive *Pygo1/2*-mutant mice and control siblings at 10.5 dpc and performed RNA-seq (n=4, Figure 6a). This analysis revealed a reproducible set of deregulated genes: among these, 65 (44 of which were downregulated and 21 upregulated) displayed high statistical significance (p-value<0.05, fold change <0,66 and >1.5) (Figure 6b). GO analysis associated a broader group of the most deregulated genes with cardiac ventricle development, embryonic limb morphogenesis and skeletal system development, recapitulating the biological processes perturbed upon *bcl9* mutation in zebrafish (Figure 6c). We confirmed the downregulation of the most relevant genes by qRT-PCR and *in situ* hybridization (*ISH*) (Figure 6d,e). Notably, the down-regulated genes comprised several genes encoding for mesodermal and NC transcription factors crucial for heart development, including *Pitx2*, *Hand2*, *Msx1*, and *Prrx1*^16,58,60,61^ (Figure 6d,e). RNA-seq of *Pygo1/2*-mutant hearts at 12.5 dpc when septation, valve formation, and myocardium proliferation and thickening have occurred^13^, showed more profound gene expression changes, with 1597 genes being deregulated (533 down, 1064 up, p-value<0.05, n=3). Consistent with progressive deterioration of cardiac development, the deregulated genes at 12.5 dpc fall into broader GO categories, including perturbed metabolic processes, protein translation, ribosome assembly, muscle formation, and increased regulation of programmed cell death (Supplementary Table 3). Several genes deregulated at 10.5 dpc, including *Msx1*, *Prrx1* and *Pitx2,* are known regulators of tissue-specific gene regulatory networks^16,61–64^, suggesting that β-catenin, via PYGO-BCL9, directly controls a subset of these genes, and that their early perturbation leads to the multitude of deregulated cellular processes at 12.5 dpf as a secondary consequence.

**Figure 6:**
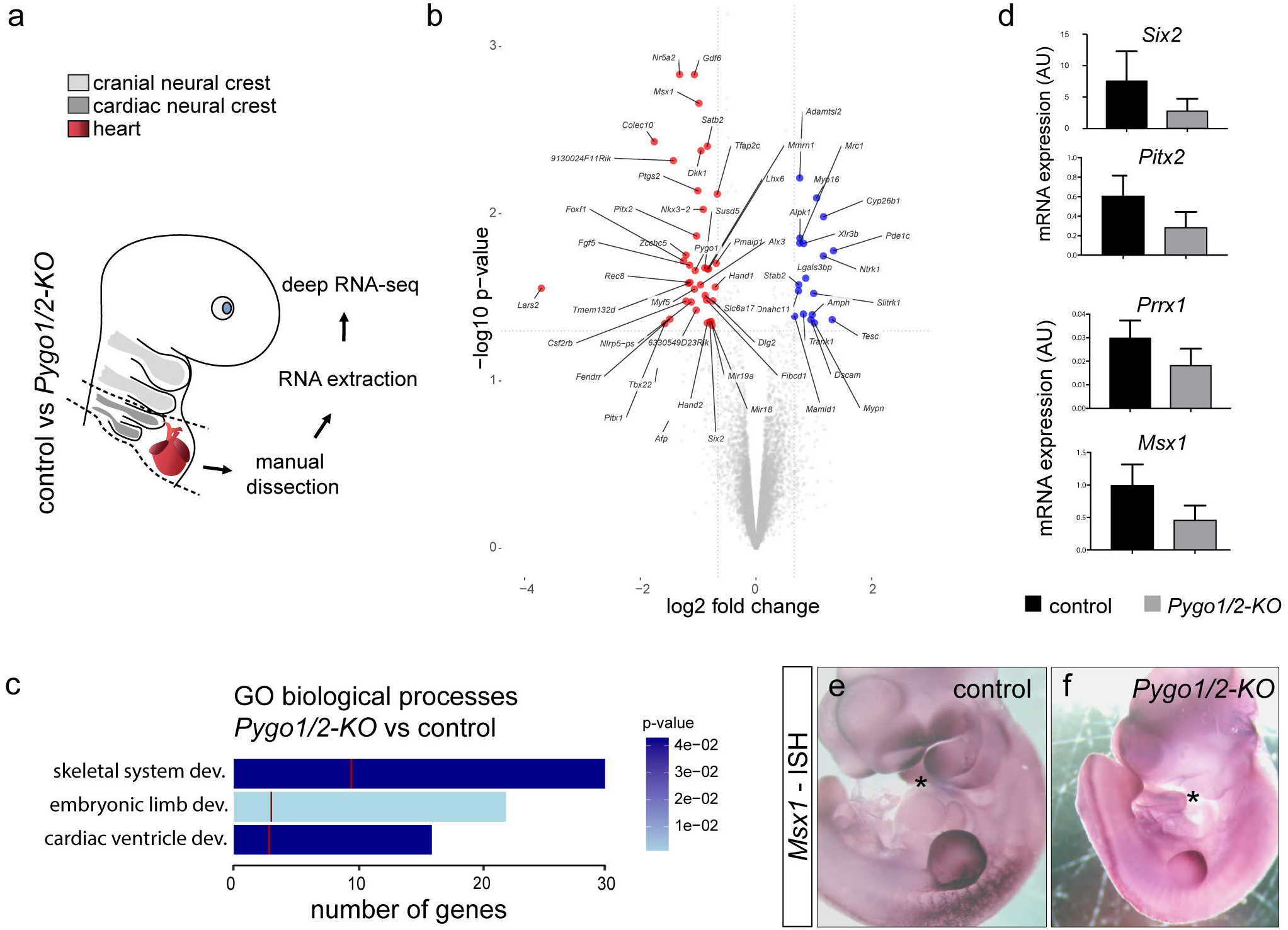
The Pygo-BCL9-β-catenin transcriptional complex controls the expression of a heart-specific genetic program. (**a**) Schematic representation of the RNA-seq approach used for *Pygo1/2-KO* mutant mouse embryos. RNA extraction was performed on manually dissected branchial arches 3-6 and hearts of 10.5 dpc embryos as indicated by the dashed outline. (**b**) Volcano plot depicting the set of upregulated (23) and downregulated (43) genes in *Pygo1/2* knockout (KO) mouse embryos with a fold change of &0.66 and &1.5, p&0.05. (**c**) Gene Ontology (GO)-based categorization of a broader group consisting of the 500 most deregulated genes (based on p-value) indicates the biological processes affected. The total bar length indicates the number of genes that categorize to the indicated GO term, the red line indicates the expected number of genes in a random deep sequencing analysis, the color of the bar indicates the p-value for the gene enrichment in our analysis. (**d**) RT-qPCR validation of a subset of target genes. mRNA expression level was calculated based on two housekeeping genes, and it is shown as arbitrary units (AU). (**e**) Whole-mount in situ hybridization (ISH) in 10.5 dpc control (left) and *Pygo1/2-KO* (right) mouse embryos shows tissue-specific (branchial arches) downregulation of the target gene *Msx1*.

To discover the genomic locations that are bound by the PYGO-BCL9-β-catenin complex, we performed chromatin immunoprecipitation followed by sequencing (ChIP-seq) for β-catenin and PYGO2 at 10.5 dpc from dissected pharyngeal arches 3-6 and the heart, as carried out for the transcriptome analysis (Figure 7a). ChIP-seq analysis assigned a total of 982 high-confidence genomic loci occupied by β-catenin, and 5252 by PYGO2 (Figure 7b); the broader chromatin association of PYGO2 is in line with the known role of these factors as broad chromatin code-readers^45,65^ and their Wnt-independent functions^28^. Nonetheless, co-occupancy of β-catenin and PYGO2 occurred at 59% of all β-catenin peaks (577/982). We focused our attention on these overlapping peaks, among which we found regions that mapped to the vicinity of canonical Wnt targets, including *Axin2* and *Lef1* (Figure 7c). These regions corresponded to previously described Wnt-regulatory-elements (WRE) at these loci found in other cell types^66,67^, and displayed the consensus binding sequence of the TCF/LEF transcription factors (Figure 7d) Therefore, while systemic canonical Wnt target genes retain most of their activity in *Pygo* and *Bcl9* mutants, these data combined with the tissue-specific loss of *Axin2* expression in *Bcl9/9l* mutants (Figure 3g-l) indicate a tissue-specific contribution from the Pygo-BCL9-β-catenin complex in branchial arches and heart cells. GO analysis of target genes that were associated to β-catenin/Pygo2 co-occupied regions (based on proximity to known transcriptional start sites (TSS)) revealed enrichment for genes involved in heart morphogenesis and valve formation, cardiac chamber septation, and skeletal development (Figure 7e,f). Intersecting the data of the RNA-seq and the ChIP-seq experiments converged on a short list of heart-associated regulatory genes that are potentially controlled by the Pygo-BCL9-β-catenin complex, including *Msx1, Prrx1*, and *Six2* (Figure 7g).

Collectively, our data support the notion that a subset of heart-specific transcriptional regulators is controlled by BCL9 and Pygo as a consequence of their context-dependent interaction with β-catenin.

**Figure 7:**
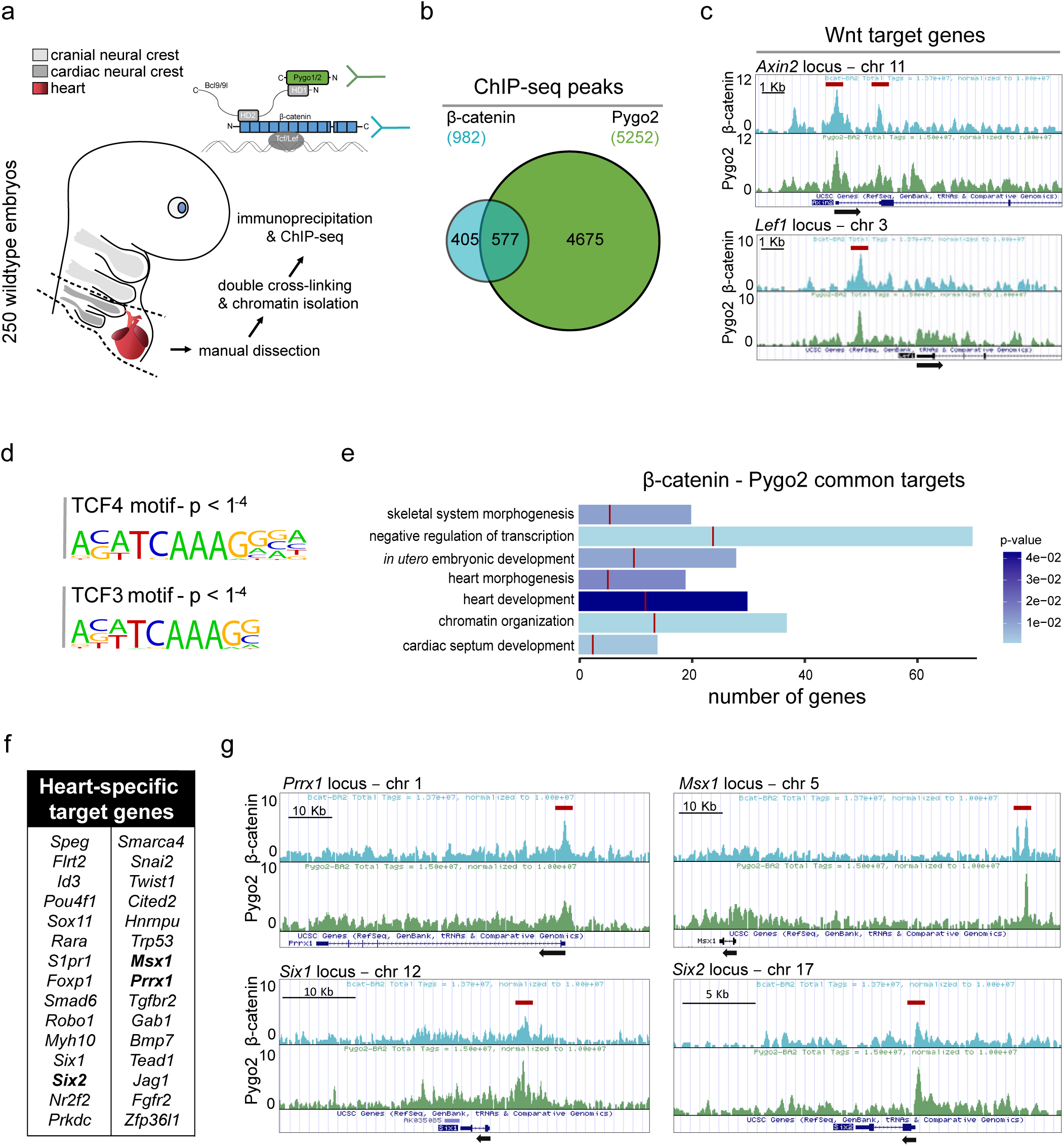
Pygo2 and β-catenin co-occupy regulatory regions of Wnt-target and heart-specific genes. (**a**) Schematic representation of the ChIP-seq approach. Branchial arches 3-6 and hearts of 10.5 dpc wildtype embryos were dissected manually as indicated by the dashed outline. Immunoprecipitation was done for β-catenin and Pygo2. (**b**) Venn diagram depicting the peak overlap between β-catenin and Pygo2 occupancy. (**c**) β-catenin and Pygo2 ChIP-seq peaks on the Wnt responsive elements of the prototypical target genes *Axin2* (upper panel) and *Lef1* (lower panel). Red bars indicate the position of previously described WREs. ChIP-seq custom tracks are visualized in the UCSC genome browser. (**d**) The binding motif of TCF/LEF transcription factors is present within β-catenin peaks. (**e**) Gene Ontology analysis of the common β-catenin and Pygo2 target genes reveals the direct regulation of processes associated with heart development. The total bar length indicates the number of genes that categorize to the indicated GO term, the red line indicates the expected number of genes in a random deep sequencing analysis, the color of the bar indicates the p-value for the gene enrichment in our analysis. (**f**) List of the 30 direct Pygo2-BCL9/L-β-catenin target genes that have been previously implicated in heart development. (**g**) β-catenin and Pygo2 ChIP peaks within the regulatory regions of genes involved in cardiac development and found downregulated upon Pygo2 loss (see Figure 6). Red bars indicate the position of previously described WREs. ChIP-seq custom tracks are visualized in the UCSC genome browser.

## Discussion

Numerous proteins have been shown to function with nuclear β-catenin, but only few are considered universal core components of the canonical Wnt pathway. Contrary to the mandatory requirement in *Drosophila*^23,65^, genetic evidence in mammals has questioned the significance and involvement of BCL9-mediated tethering of the histone reader Pygo to β-catenin in Wnt target gene control^29–31^. Here, by combining *in vivo* analyses in zebrafish and mouse, we revealed a conserved contribution of BCL9 and Pygo proteins as context-dependent regulators of Wnt/β-catenin signaling in vertebrates, in particular during heart development. Our *in vivo* data clarifies, consolidates, and extends previous work in several models on BCL9 and Pygo protein function. Additionally, our results implicate perturbed BCL9/9L levels as potentially causative for human CHDs, as it occurs in copy number variants of *1q21.1*.

Biochemical and cell culture-based evidence has linked the BCL9-Pygo complex to a variety of nuclear processes independent of β-catenin and TCF. In parallel, *in vivo* evidence for BCL9 and Pygo function has suggested divergent functions for these proteins in mammals versus fishes and amphibians: while overexpression and morpholino-based experiments in zebrafish and *Xenopus* have assigned β-catenin-dependent functions for BCL9 and Pygo proteins already during gastrulation, loss-of-function studies during mouse development have uncovered mainly Wnt-independent phenotypes^25,26,29^. To date, no clear consensus of their function during vertebrate development has emerged. By using genetic and chemical loss-of-function studies both in zebrafish and mouse, we define the *in vivo* requirement of BCL9 and Pygo during vertebrate development, and specifically their role in relation to β-catenin. The phenotypes resulting from abrogating their function imply that the Pygo-BCL9-β-catenin complex, while being dispensable during early embryogenesis, is fundamental as context-dependent transcriptional regulator during several tissue-specific processes, in particular during heart development.

We corroborate this conclusion by several lines of evidence. Previous work and our results confirm that various genetic loss-of-function permutations of BCL9 and PYGO proteins do not perturb gastrulation in mouse^29^ (Figures 2 and 3). In zebrafish, using CRISPR-Cas9, we generated mutants with a premature stop codon between the HD1 and HD2 coding sequence of *bcl9* and *bcl9l*; these mutations should uncouple BCL9/9L-PYGO1/2 from engaging with β-catenin. Zebrafish Bcl9 and Bcl9l are maternally contributed, yet we failed to detect any gastrulation defects previously reported for morpholino knockdown of *Xenopus XBcl9* and zebrafish *bcl9l*^25,26^. We substantiated this conclusions in three experimental setups: i) both zygotic and maternal-zygotic *bcl9l^Δ4/Δ4^* homozygous mutants are viable and fertile (Supplementary Figure 1); ii) selective chemical interruption of the BCL9/9L-β-catenin interaction during gastrulation or later phenocopies the *bcl9^Δ29^*-mutant phenotype (Figure 4); iii) chemical perturbation of the BCL9/9L-β-catenin interaction in *bcl9* heterozygous mutant crosses increases penetrance of strong phenotypes presumably due to a more sensitive genetic background of heterozygotes but never leads to a stronger expressivity or gastrulation defects (Supplementary Figure 8). A potential explanation for the discrepancy with morpholino-based findings could be that our zebrafish alleles are not null alleles and residual Bcl9/9l products possibly rescue the phenotype. We further cannot rule out genetic compensation during gastrulation for *bcl9* by *bcl9l* and *vice versa*^41^, despite absence of such an effect at later stages in our double-mutants (Figure 1). Nonetheless, the genetic evidence together with chemical perturbation of the Bcl9/9l-HD2 by LH-4-20^49^ support only a tissue-specific contribution of BCL9-tethered Pygo to β-catenin-based Wnt signaling in zebrafish development.

Both zebrafish and mouse embryos with perturbed BCL9 and Pygo broadly maintain transcriptional activity of β-catenin (Figures 3 and 4). Nonetheless, following our observation of cardiac defects in interaction-perturbing *Bcl9* and *Pygo* mutants, transcriptome analysis (Figure 6) and chromatin occupancy of β-catenin and PYGO (Figure 7) uncovered a collection of cardiac and pharyngeal genes sensitive to disruption of the PYGO-BCL9-β-catenin complex. Among these, we found common pan-β-catenin target genes, including *Axin2* and *Lef1*. Importantly, however, we predominantly uncover genes associated with tissue-specific functions (Figures 7). Our data support models of canonical Wnt target gene expression by context-dependent nuclear β-catenin interactions, and define BCL9 and Pygo proteins as selectively required β-catenin interactors in vertebrate heart development. Of note, cardiac defects in mice such as AV valve malformations lead to embryonic lethality between 13.5 and 16.5 dpc^68,69^, suggesting that the cardiac malformations could indeed cause the lethality of Pygo1/2-mutant and Bcl9/9l-ΔHD1 embryos. The limb and additional skeletal phenotypes observed in our mutants deserve detailed future analysis. Our findings further emphasize that BCL9 and Pygo cannot be strictly defined as core components of canonical Wnt pathway; if their mandatory contribution to canonical Wnt signaling in *Drosophila* represents an ancestral function or a specialization warrants further investigation.

Multiple contributions of Wnt signaling during vertebrate heart formation have previously been uncovered: waves of activation and inhibition of Wnt/β-catenin signaling are essential for the balance between proliferation versus differentiation of cardiac progenitors^17–19,70,71^, for endocardial invagination and EMT during valve formation, and in and neural crest cell migration into the heart^15,46,72–74^. Consequently, the alteration of canonical Wnt signaling can affect several aspects of heart formation. The cardiac defects caused by mutations in *Bcl9* and *Pygo* recapitulate previous perturbations of canonical Wnt signaling obtained by overexpression of the extracellular Wnt antagonist Dkk1 at discrete time points during cardiac development^46^. In this context, BCL9 and Pygo contribute to orchestrating the downstream expression of selected mesodermal heart and neural crest regulators, including *Msx1, Prrx1,* and *Six2*, as further emphasized by β-catenin and PYGO2 binding in the vicinity of these loci (Figure 7g). Of note, individual deregulation of each of these factors can cause specific developmental heart defects: *Msx1, Prrx1,* and *Six2* were previously implicated in CNC cells proliferation, in SHF-dependent OFT formation, and in EMT during AV valves formation, respectively^58,61,75^. Their downregulation, in turn, causes the subsequent deregulation of additional critical factors for heart morphogenesis, including *Hand2, Pitx2,* as well as members of the FGF signaling cascade (Supplementary Table 3)^76,77^. Notably, *Pitx2*, a seemingly secondary target of Wnt signaling in this context, is involved in septation and left-right patterning of the heart, as well as in modulation of a Tbx5-dependent GRN in heart homeostasis^64^. Loss-of-function of *Hand2* and *Pitx2* is individually associated with several heart defects reminiscent of the phenotypes caused by perturbation of the PYGO-BCL9-β-catenin complex^4,5,13,16^. These findings indicate that canonical Wnt/β-catenin signaling modulates a particular cardiac GRN via the direct activation of transcription factor genes including *Msx1, Prrx1, Six2*, and possibly *Hand2* using the context-specific β-catenin co-factors BCL9 and PYGO.

An imbalance of BCL9 and Pygo levels both by loss and by overexpression affects β-catenin-dependent canonical Wnt readouts in several systems. In *Drosophila*, misexpression of a *lgs* variant that fails to bind β-catenin/Armadillo impairs canonical Wnt signaling, as does overexpression of BCL9 in human cell culture^22,78^. Altering the level of BCL9 in *Xenopus* leads to gastrulation defects that are reminiscent of defects in β-catenin function during organizer formation^26^. These effects can be explained by sequestration of Pygo and β-catenin by elevated BCL9 levels, resulting in a net loss of functional Pygo-BCL9-β-catenin complexes akin to reduced or lost BCL9. In non-syndromic, sporadic cases of human CHD (with or without additional extra-cardiac phenotypes), copy number gains and losses have repeatedly been found in the genomic region *1q21.1* that includes the *BCL9* locus^32–38^. CNVs that lead to either loss or gain of *BCL9* copies could perturb the sensitive balance of the PYGO-interacting co-activator function of BCL9 with β-catenin during human heart development. Furthermore, mutations in *BCL9L*, including changes in the HD1, have been genetically linked in a small pedigree to heterotaxia (HTX) with congenital cardiac malformations including ventricular and atrial septal defects^39^. Our genetic and functional data reveal a causative connection between varying Wnt/β-catenin signaling levels by mutations in *Bcl9* and *Pygo* with developmental heart defects in both zebrafish and in mice; therefore, our findings indicate that CNVs of, or mutations in, *BCL9* and *BCL9L* contribute to the etiology of human CHDs through tissue-specific perturbation of canonical Wnt signaling.

## Acknowledgements

We thank Sibylle Burger, Seraina Bötschi, and Eliane Escher for technical support in zebrafish husbandry and mouse genotyping, Dr. Stephan Neuhauss for zebrafish support, Dr. Haitao Ji for chemical compounds, the ZBM at UZH for imaging support, and the members of the Mosimann and Basler labs for constructive input. This work has been supported by a Swiss National Science Foundation (SNSF) professorship [PP00P3_139093] and SNSF R’Equip grant 150838 (Lightsheet Fluorescence Microscopy), a Marie Curie Career Integration Grant from the European Commission [CIG PCIG14-GA-2013-631984], the Canton of Zürich, the UZH Foundation for Research in Science and the Humanities, and the Swiss Heart Foundation to C.M.; by the Swiss National Science Foundation (SNF) to K.B. and grants from the Forschungskredit of the University of Zürich to C.C.; J.R. acknowledges support from MINECO FIS2016-77892-R.

## Author contribution

C.C., A.F. and D.Z. designed and performed the experiments, interpreted the data and wrote the manuscript; E.C. and L.K. assisted with zebrafish mutants generation and characterization; E.M.C. performed bioinformatics analyses; T.V. assisted with mouse breeding and genotyping; G.H. helped with the design of the work and critically revised the article; J.R. performed and analyzed the mouse SPIM images; N.V. and M.A. performed the initial analyses on *Pygo1/2*-mutant mice; K.B. and C.M. supervised and assisted the research teams, interpreted results, and wrote the manuscript.

## Methods

### Zebrafish husbandry and transgenic strains

All zebrafish embryos were raised and maintained in E3 medium at 28.5°C without light cycle essentially as previously described^79^. All experiments were performed on embryos and larvae up to 5 dpf and older larvae only kept for raising mating pairs in agreement with procedures mandated by the veterinary office of UZH and the Canton of Zürich. Embryo staging was done according to morphological characteristics corresponding to hours post fertilization (hpf) or days post fertilization (dpf) as described previously^80^. Previously established transgenic zebrafish lines used for this study include *Tg(-6.35drl:EGFP)*^81^ and *Tg(7xTCF-Xla.Siam:nlsmCherry)^ia5^*^48^.

### CRISPR-Cas9 mutagenesis

CRISPR-Cas9 mutagenesis was essentially performed as described in Burger et al. 2016^82^. Oligo-based sgRNA templates^83^ were generated by PCR amplification using the invariant reverse primer *5’-AAAAGCACCGACTCGGTGCCACTTTTTCAAGTTGATAACGGACTAGC CTT-3’* and forward primers of the sequence *5’-GAAATATTTAGGTGACACTATA-(N20-22)-GTTTTAGAGCTAGAAATAGC-3’* with N representing the 20 nucleotides of the *sgRNA* target sequence plus up to 2 *G*s at the 5’-end for successful T7 *in vitro* transcription. sgRNA (plus added 5’Gs in brackets) sequences were: i) *bcl9 5’-(G)GCCTGACTGCGCTACTTTAG-3’*; ii) *bcl9l 5’-(G)GATTTAGGTGTGCCAATCGG-3’*; iii) *pygo1 5’-GGACTTCCCAGTAGCAGCCC-3’*; iv) *pygo2 5’-(G)GCCGATGGTTGACCACCTGG-3’*. Crispants were raised to adulthood and crossed to wildtypes to make F1 germline mutants. All analyses and experiments were taken out on F2 mutant generations and beyond. Genotyping primers were designed to either amplify target regions of mutated alleles that were subsequently analyzed via sequencing (for *pygo1Δ1*) or high-percentage gel electrophoresis (for *bcl9^Δ29^*) or allele-specific primers were designed to bind to mutated vs. wildtype sequences specifically (for *bcl9l^Δ4^* and *pygo1*^*Δ5*^). Primers used were: i) *bcl9 5’-GGTGGAAAGCCCCAACTCC-3’* (fwd), *5’-CGTGTGCCAACTGCTGGTGG-3’* (rev); ii) *bcl9l 5’- CACTTGCAGGTGCTGCATGG-3’* (fwd), *5’- CTTTGAGATTTAGGTGTGCCGG-3’* (rev); iii) *pygo1 5’-CACTTTTACTGACCCCCACAC-3’* (fwd), *5’-GGACTTCCCAGTAGCAGGA-3’* (rev); iv) *pygo2 5’-GCCCAGAGAAAAAGAAGAGG-3’* (fwd), *5’- GCTGTCCACTTCCAGGTCC-3’* (rev). Genotyping results were analyzed and alleles visualized using CrispRVariantsLite^84^.

### Chemical treatments

Wildtypes, *Tg(7xTCF-Xla.Siam:nlsmCherry)*^*ia5*^, and embryos obtained from *bcl9^Δ29^* heterozygous incrosses were treated with LH-2-40 to globally perturb Bcl9ΔHD2-beta-catenin interaction at the respective developmental stages. LH-2-40 was kindly received from the laboratory of Dr. Haitao Ji (Moffitt Cancer Center, Tampa/FL, USA). Single-use LH-2-40 stocks were kept at a concentration of 100 mM in DMSO at -80°C and thawed and diluted in E3 to a working concentration indicated in individual experiments directly before administration to the embryos.

### Alcian Blue staining

Wildtype, *bcl9*, and *pygo1/2* mutant larvae were fixed in 4% Paraformaldehyde (PFA) overnight at 4°C and after washing in 0.1% PBS-Tween (PBST) stained in Alcian blue staining solution (0.1 g Alcian blue, 70 mL ethanol, 30 mLl glacial acetic acid) overnight at room temperature. Embryos were washed in Ethanol and transferred through an Ethanol series to PBST and subsequently bleached in hydrogen peroxide (3% H2O2 in 1% KOH in PBS) for 1 hour or until pigments of specimens became transparent.

### Whole-mount in situ hybridization

First-strand complementary DNA (cDNA) was generated from pooled zebrafish RNA isolated from different developmental stages using Superscript III First-Strand Synthesis kit (Invitrogen). DNA templates were generated using first-strand cDNA as PCR template and following primers *5’-TTACGTATGCAGCCTTCTCG-3’* and *5’-GGTTCATGGGGTAACTGTGG-3’ (vcana*), *5’-ACGGATCAAGTAAGCGAAGG-3’ and 5’-GTTCCTCCAGTTCGTTTTGC-3’ (myh6), 5’-GAGCTGCGTCTTACGAGTCC-3’ and CAGACTGGCTCTCCTTCTGC-3’* (*gata4*), *5’-ACTGATGAGGACGAGGAAGG-3’ and 5’-GACTCGGAATCCTTCAGTGG-3’ (spry4)*. For *in vitro* transcription initiation, the T7 RNA polymerase promoter *5'-TAATACGACTCACTATAGGG-3’* was added to the 5’-end of reverse primers. The DNA template for *nkx2.5* was amplified from a plasmid kindly contributed by the lab of Dr. Daniela Panáková (Max-Delbrück Center for Molecular Medicine, Berlin-Buch, Germany) using *T3* and *T7* primers. PCR reactions were performed under standard conditions using Phusion High-Fidelity DNA Polymerase (ThermoFisher Scientific). RNA probes were generated via overnight incubation at 37°C using T7 RNA polymerase (20 U/µl) (Roche) and Digoxigenin (DIG)-labeled dNTPs (Roche). The resulting RNA was precipitated in lithium chloride and Ethanol.

Embryos were fixed in 4% PFA overnight at 4°C, transferred into 100% methanol and stored at -20°C. ISH of whole mount zebrafish embryos was performed essentially according to standard protocols^85^.

### Microscopy and data processing

Brightfield (BF), basic fluorescence, and ISH imaging was performed using a Leica M205FA equipped with a DFC450 C camera. Detailed fluorescent embryo imaging of *drl*:EGFP-expressing *bcl9* mutants was performed by Single Plane Illumination Microscopy (SPIM) with a Zeiss Lightsheet Z.1 microscope. Prior to imaging, embryos were embedded in a rod of 1% low melting agarose in E3 with 0.016% Ethyl 3-aminobenzoate methanesulfonate salt (Tricaine, Sigma) in a 50 µL glass capillary.

Image processing was done with Leica LAS, Zeiss Zen Black, ImageJ/Fiji and Adobe Photoshop and Illustrator CS6 according to image-preserving guidelines to ensure unbiased editing of the acquired image data. Quantitative data analysis was performed using GraphPad Prism 5.0. Data are presented as mean ± SEM. A lower case “n” denotes the number of embryos, while a capital “N” signifies the number of replicates. To perform SPIM on embryonic mouse heart, tissue samples were extracted at 13.5 dpc, embedded in low melting agarose, dehydrated, and cleared using Benzyl Alcohol/ Benzyl Benzoate (1:2). 3D optical sectioning of the anatomy of the samples was performed with a Qls-Scope light sheet microscopy system (Planelight S.L., Madrid, Spain) by measuring autofluorescence with a 5x objective where signal was optimal, in this case exciting at 532 nm and collecting emission at 590 +/- 40 nm. Samples where imaged from two opposing views and combined to form a single volumetric image with an isotropic voxel size of 2.6 μm.

### Mouse lines

Knock-in mutants in *Bcl9* and *Bcl9l* were generated by standard techniques, as previously described^29^. Briefly, the targeting vector was electroporated into *BA1* (*C57BL/6* × *129/SvEv*) hybrid embryonic stem cells. After selection with the antibiotic G418, surviving clones were expanded for PCR and Southern blotting analyses to confirm recombinant embryonic stem cell clones. Mouse embryonic stem cells harboring the knock-in allele were microinjected into *C57BL/6* blastocysts. Resulting chimeras were bred to wildtype *C57BL/6N* mice to generate F1 heterozygous offspring. Neo cassette excision was obtained by crossing heterozygous knock-in animals with mice expressing Flp recombinase. All mouse experiments were performed in accordance with Swiss guidelines and approved by the Veterinarian Office of the Kanton of Zurich, Switzerland.

### Histological analysis

Embryos between day 9.5 and 14.5 dpc were fixed overnight in 4% paraformaldehyde at 4°C, dehydrated and embedded in paraffin. Sections were stained with haematoxylin and eosin for histological analysis. The same paraffin embedded material was sectioned under RNase free conditions for in situ hybridization.

### Mouse *in situ* hybridization

Whole-mount *in situ* hybridization (ISH) was performed as described previously^86^. Digoxigenin-labeled probes (Roche) were detected by enzymatic color reaction using alkaline phosphatase-conjugated anti-digoxigenin Fab fragments (1:1000, Roche) and BM purple alkaline phosphatase substrate (Roche). Digoxigenin-labeled RNA probes were detected with peroxidase-conjugated anti-digoxigenin Fab fragments (1:500, Roche) followed by fluorescence detection using Tyramide Signal Amplification (PerkinElmer). Mouse antisense RNA probes were as described in supplementary materials. Quantitative RNA in situ hybridization was performed using RNAscope 2.5 (ACD) according to manufacturers’ instruction. Mean intensity measurements were performed using ImageJ.

### Intracardiac injection of India ink

For the analysis of the cardiovascular system, india ink was injected into the left ventricle of mouse embryos at 14.5 dpc, using a finely drawn glass pipette, right after embryonic dissection when the heart was still beating. The embryos were subsequently fixed in 4% paraformaldehyde and hearts were dissected for the anatomic examination.

### RNA sequencing

*Zebrafish samples (polyA selection):* Zebrafish embryos from *bcl9^Δ29/wt^* crosses were dissected at 54 hpf: a cut was made between the anterior part of the embryo (containing the head, pharyngeal and cardiac structures) and the posterior part starting from the beginning of the yolk extension. The anterior tissues were directly snap-frozen in liquid nitrogen in single tubes of PCR 8-tube strips. The posterior trunks and tails were transferred in 50 mM NaOH and genomic DNA was extracted with alkaline lysis. Single embryos were genotypes using the target sequence primers listed above and PCR products were separated through high-percentage gel electrophoresis leading to a separation of the wildtype allele and mutant allele with the following outcome: i) a low running band in *bcl9^Δ29^* mutants; ii) a high running band in wildtypes; iii) two bands in *bcl9^Δ29/wt^* heterozygous mutants. All snap-frozen anterior parts of *bcl9^Δ29^* mutants and of wildtypes were pooled in two separate tubes and RNA isolation performed with the RNeasy Plus Mini Kit (Qiagen). The whole procedure was repeated for a total of three clutches making three independent replicates.

The TruSeq mRNA stranded kit from Illumina was used for the library preparation with 250 ng of total RNA as input. The libraries were 50-bases sequenced on an Illumina HiSeq 2500 sequencer. The quality control of the resulting reads was done with FastQC and the reads mapped to the UCSC *Danio rerio* danRer10 genome with the TopHat v.2.0.11 software. For differential expression analysis the gene features were counted with HTSeq v.0.6.1 (htseq-count) on the UCSC danRer10 gene annotation. The normalization and differential expression analysis were performed with R/Bioconductor package EdgeR v. 3.14. The p-values of the differentially expressed genes are corrected for multiple testing error with a 5 % false discovery rate (FDR) using Benjamini-Hochberg (BH). The volcano plot was done selecting the genes that have > 10 counts per million (CPM) in at least 3 samples, p-value < 0.12 and absolute value of log fold change (logFC) above 1. STRING analysis was performed on differentially expressed genes implicated in cardiac development in the literature (see Tables S1,2). Permalink to analysis: http://bit.ly/2wmba7z.

*Mouse samples (ribo depletion):* The Illumina TruSeq stranded Total RNA library Prep kit with RiboZero was used for the library preparation with 300 ng of total RNA as input. The libraries were 100-bases sequenced on an Illumina HiSeq 4000 sequencer. The quality control of the resulting reads was done with FastQC and the reads mapped to the UCSC *Mus musculus* mm10 genome with the TopHat v.2.0.11 software. For differential expression analysis the gene features were counted with HTSeq v.0.6.1 (htseq-count) on the UCSC mm10 gene annotation. The normalization and differential expression analysis were performed with R/Bioconductor package EdgeR v. 3.14. The p-values of the differentially expressed genes are corrected for multiple testing error with a 5 % false discovery rate (FDR) using Benjamini-Hochberg (BH).

### qRT-PCR

Quantitative real-time SYBR Green-based PCR reactions were performed in triplicate and monitored with the ABI Prism 7900HT system (Applied Biosystem).

### Chromatin Immunoprecipitation (ChIP)

Ca. 200 branchial arches of E10.5 embryos were dissected and dissociated to a single cell suspension. Cells were cross-linked in 20 ml PBS for 40 min with the addition of 1.5 mM ethylene glycol-bis(succinimidyl succinate) (Thermo Scientific, Waltham, MA, USA), for protein-protein cross-linking, and 1% formaldehyde for the last 20 min of incubation, to preserve DNA-protein interactions. The reaction was blocked with glycine and the cells were subsequently lysed in 1 ml hepes buffer (0.3% SDS, 1% Triton-X 100, 0.15 M NaCl, 1 mM EDTA, 0.5 mM EGTA, 20 mM HEPES). Chromatin was sheared using Covaris S2 (Covaris, Woburn, MA, USA) for 8 min with the following set up: duty cycle: max, intensity: max, cycles/burst: max, mode: Power Tracking. The sonicated chromatin was diluted to 0.15% SDS and incubated overnight at 4°C with anti IgG (Santa Cruz), anti-Pygo2 (Novus Biological, NBP1-46171) and anti–β-catenin (Santacruz, sc-7199). The beads were washed at 4°C with wash buffer 1 (0.1% SDS, 0.1% deoxycholate, 1% Triton X-100, 0.15 M NaCl, 1 mM EDTA, 0.5 mM EGTA, 20 mM HEPES), wash buffer 2 (0.1% SDS, 0.1% sodium deoxycho– late, 1% Triton X-100, 0.5 M NaCl, 1 mM EDTA, 0.5 mM EGTA, 20 mM HEPES), wash buffer 3 (0.25 M LiCl, 0.5% sodium deoxycholate, 0.5% NP-40, 1 mM EDTA, 0.5 mM EGTA, 20 mM HEPES), and finally twice with Tris EDTA buffer. The chromatin was eluted with 1% SDS, 0.1 M NaHCO3, de-crosslinked by incubation at 65°C for 5 h with 200 mM NaCl, extracted with phenol-chloroform, and ethanol precipitated. The immunoprecipitated DNA was used as input material for DNA deep sequencing.

*Data analysis – peak calling*: All sequenced reads were mapped using the tool for fast and sensitive reads alignment, Bowtie 2 (http://bowtie-bio.sourceforge.net/bowtie2/index.shtml), onto the UCSC mm10 reference mouse genome. The command "findPeaks" from the HOMER tool package (http://homer.salk.edu/homer/) was used to identify enriched regions in the beta-catenin immunoprecipitation samples using the "-style = factor" option (routinely used for transcription factors with the aim of identifying the precise location of DNA-protein contact). Input and IgG samples were used as enrichment-normalization controls. Peak calling parameters were adjusted as following: L = 4 (filtering based on local signal), F = 4 (fold-change in target experiment over input control). Annotation of peaks' position (i.e. the association of individual peaks to nearby annotated genes) was obtained by the all-in-one program called "annotatePeaks.pl". Finally, the HOMER command "makeUCSCfile" was used to produce bedGraph formatted files that can be uploaded as custom tracks and visualized in the UCSC genome browser (http://genome.ucsc.edu/).

### Immunofluorescence

FFPE sections were blocked with 5% hings, % BSA and 0.1% Tween in PBS and incubated overnight at 4°C with the following antibodies: mouse anti p53 5E2 (NovusBio), rabbit anti Sox9 (Millipore), Troponin type2 (novus biologicals), Acta2, GFP (Aves). Slides were then incubated with a fluorescently labeled secondary antibody (Alexa 488 goat anti-mouse or Alexa 555 goat anti-rabbit, Alexa 594 goat anti-chicken; 1:500). Nuclei were stained with DAPI (1:1000; Sigma).

### Branchial arch explants

Pharyngeal apparatus was dissected in PBS and resuspended in digestion mix (0.5mg collagenase/ml, 0.1% Trypsin), incubated at 37 °C for 15min. Tissue digestion was inactivated with 750ul of DMEM 10% FCS and cells were plated in fibronectin coated wells in medium according to Zhao et al., 2006^87^.

### Accession codes

ChIP-seq and RNA-seq reads are being deposited in the Gene Expression Omnibus (GEO) repository.

## Supplementary Material

**Supplementary Figure 1:**
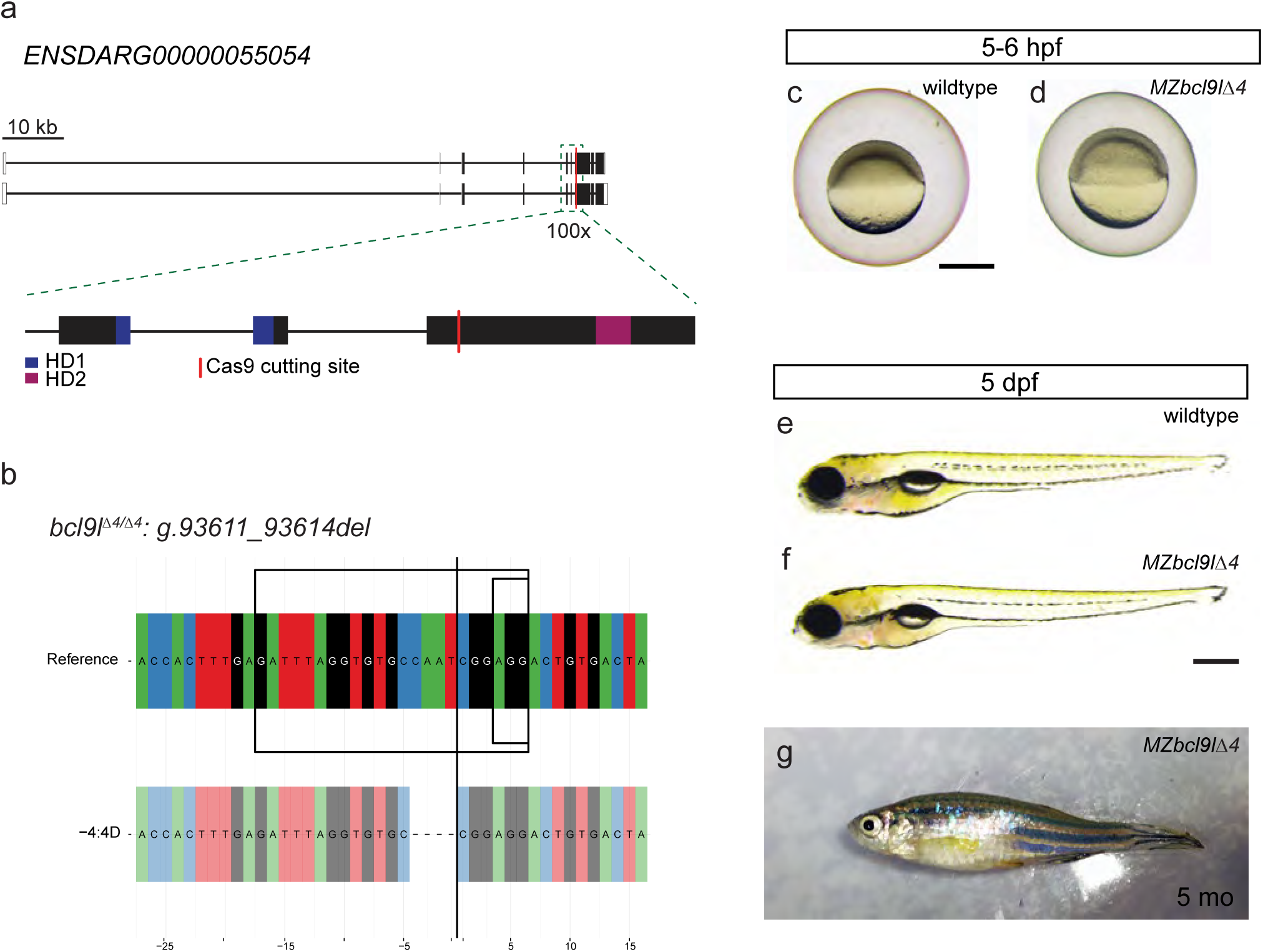
*MZbcl9l^Δ4^* mutants are viable and fertile. (**a,b**) Schematics showing the mutation induced in *bcl9lΔ4* mutants. Analogous to *bcl9* mutants (see Figure 1b,c), we designed a sgRNA that binds between the HD1 and HD2 domain. Gene locus represented as per genome annotation *Zv10* (**a**) with two isoforms that differ in the first coding exon and the untranslated regions (UTRs). The green dotted box represents a zoomed region of the gene locus, with the red line representing the location of the sgRNA used for mutagenesis; black boxes mark coding exons (CDS), white boxes mark UTRs, the blue boxes represent the part of the CDS that will contribute to HD1 and purple boxes to HD2. A 4 pb deletion at the Cas9 cutting side (**b**) leads to a frame-shift allele with a premature *STOP* codon before HD2. (c-g) Brightfield images of homozygous *bcl9l^Δ4^* mutants at different stages from gastrulation to adulthood, lateral views, anterior to the left. Maternal zygotic *bcl9l* mutants (MZbcl9^lΔ4^) mutants did not show any gastrulation defects (**c,d**) in contrast to *bcl9l* morphant phenotypes reported in the literature. At 5 dpf we could not detect any cardiac and craniofacial phenotypes (**e,f**) as observed in zygotic *bcl9^Δ29^* mutants (see Figure 1). (**g**) Representative image of a five-month old (5 mo) *MZbcl9l^Δ4^* F4 mutant obtained from a cross of two adult homozygous *bcl9l^Δ4^* mutants. Scale bars, 500 µm.

**Supplementary Figure 2:**
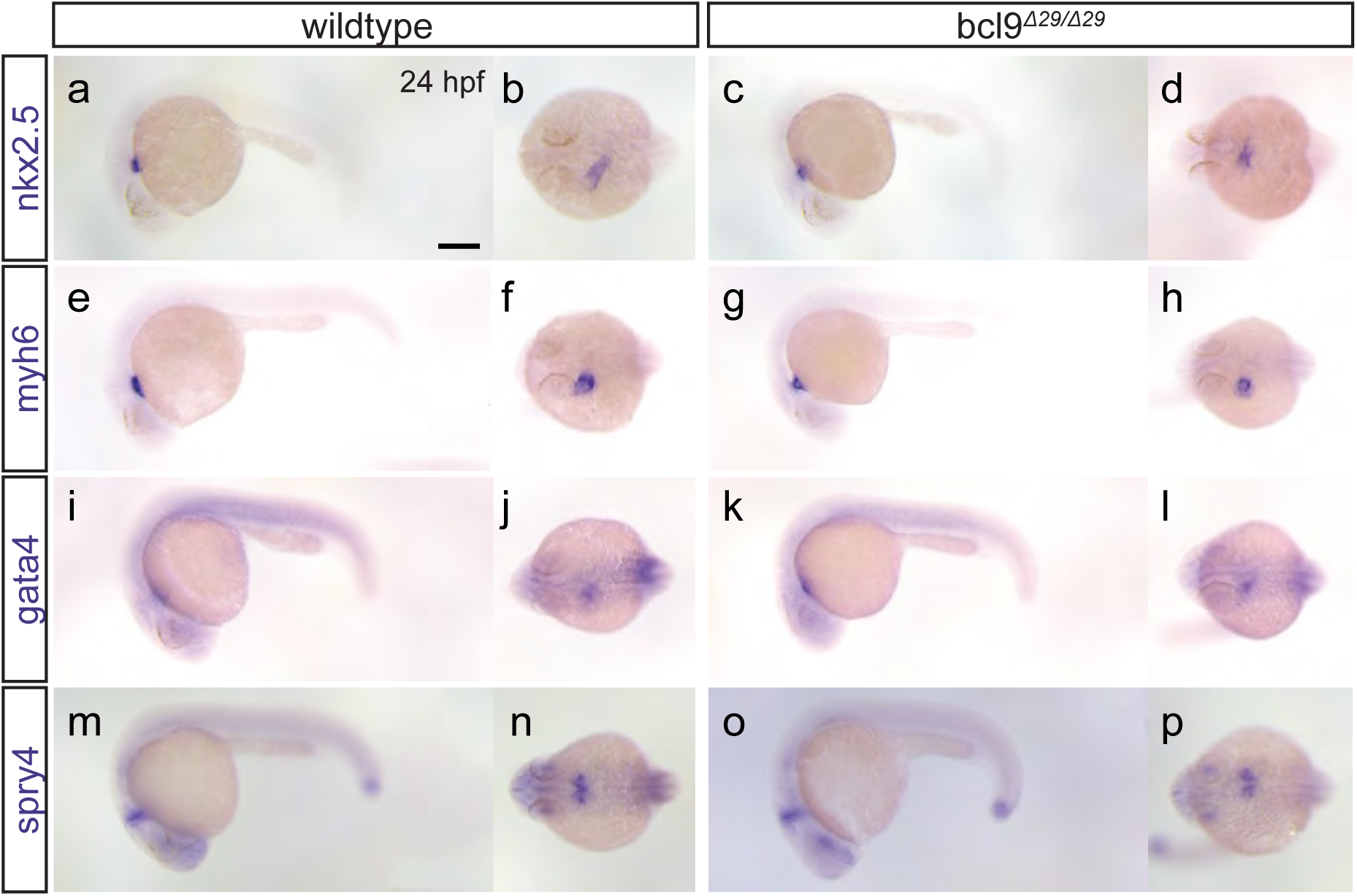
Early cardiac patterning is unperturbed in *bcl9* mutants. (**a-p**) *bcl9* mutants displayed unchanged *nkx2.5* (**a-d**), *myh6* (**e-h**), *gata4* (**i-l**), and *spry4* (**m-p**) expression at 24 hpf compared to wildtype siblings, lateral and dorsal views, anterior to the left. Scale bar, 250 µm.

**Supplementary Figure 3:**
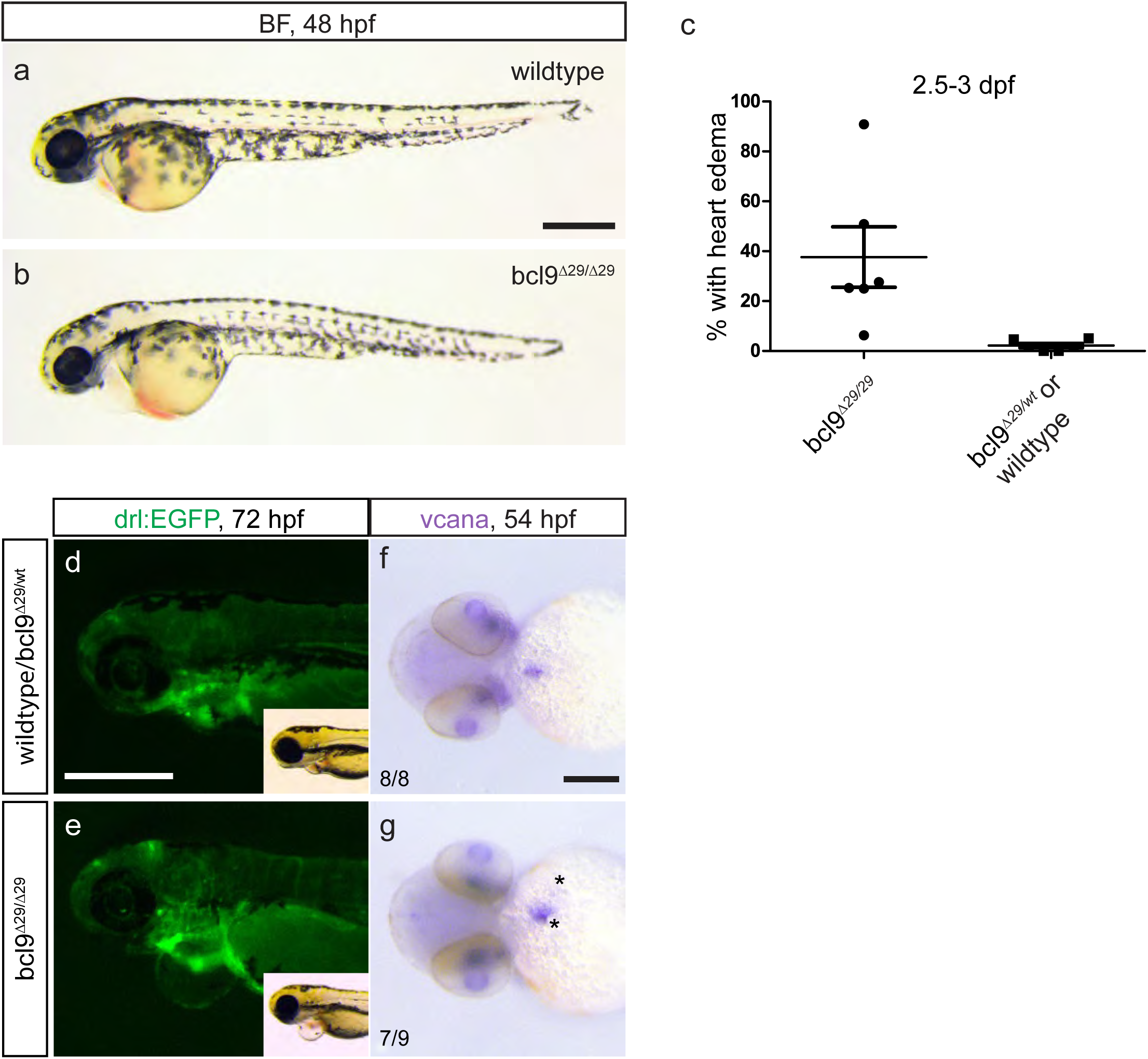
Variable cardiac phenotypes in *bcl9^Δ29^* mutants between 2-3 dpf. (**a-c**) At 48 hpf, *bcl9^Δ29^* develop pericardiac edema with a variable penetrance, lateral views (**a,b**), anterior to the left. (d,e) Fluorescent and brightfield (inlets) images of *drl*:EGFP transgenic wildtype and *bcl9^Δ29^* mutant embryos at 72 hpf, lateral views, anterior to the left. The hearts of *bcl9^Δ29^* are patterned into atria and ventricle. The two chambers are, however, miss-aligned revealing a looping defect caused by the *bcl9* mutation. (**f,g**) Whole mount ISH for *vcana* reveals expanded expression around the atrio-ventricular canal and in the atrium (asterisks) suggesting a defect in valve formation, ventral views, anterior to the left. Scale bars, 200 µm (**f,g**), 500 µm (**a,d**).

**Supplementary Figure 4:**
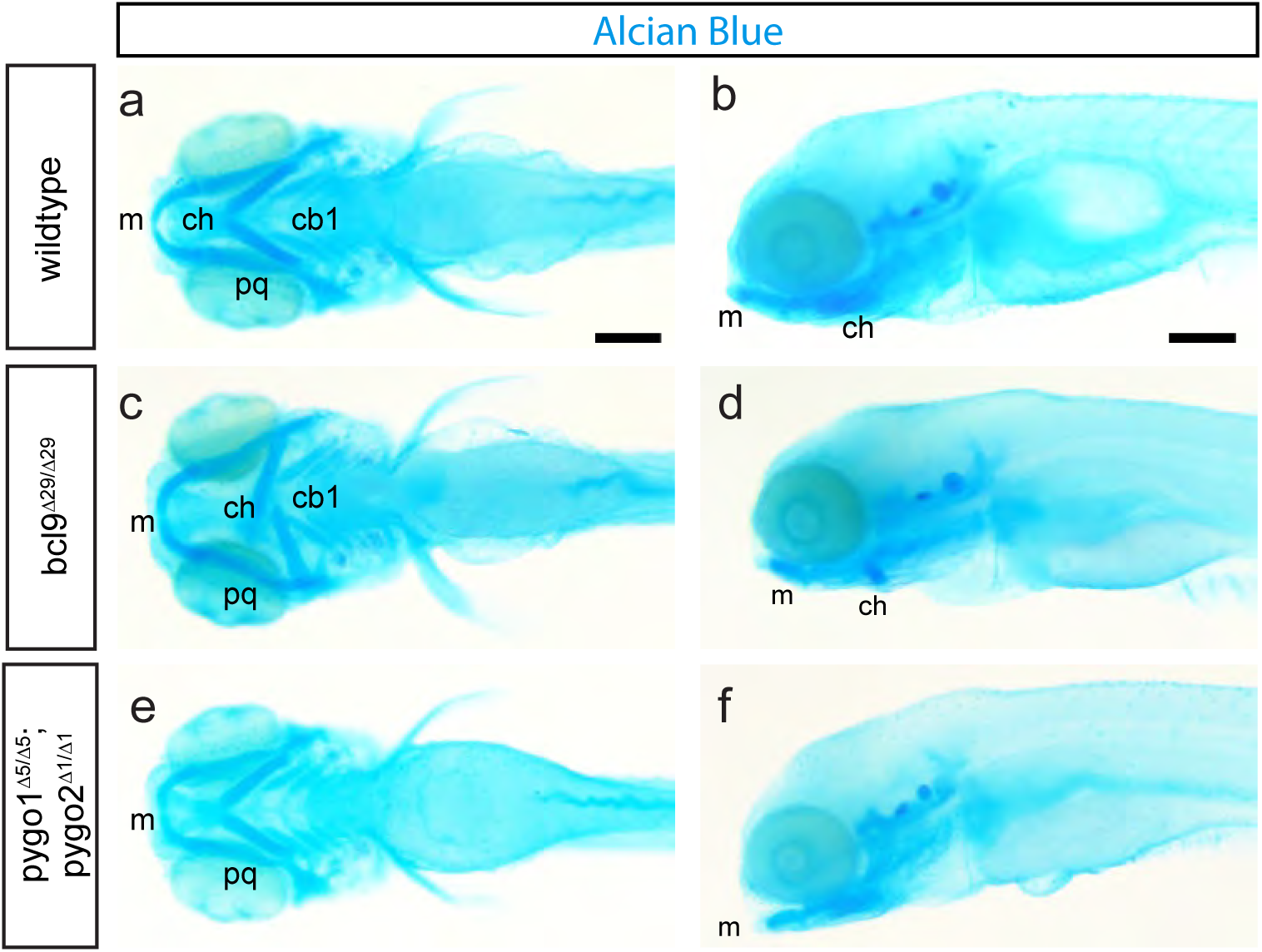
Craniofacial defects in *bcl9* and *pygo1/2* mutants. (**a-f**) Alcian blue staining of the pharyngeal cartilage of 5 dpf wildtype, *bcl9^Δ29^*, and double homozygous *pygo1^Δ5^*;*pygo2^Δ1^* larvae shown in ventral (**a,c,e**) and lateral (**b,d,f**) views, anterior to the left. *bcl9* and *pygo1/2* mutants have severe malformations of the pharyngeal apparatus with fusions defect of the ceratohyal (ch) and ceratobranchial 1 (cb1) arches and miss-shaped Meckel’s (m) and palatoquadrate (pq) cartilage. Scale bars, 100 µm.

**Supplementary Figure 5:**
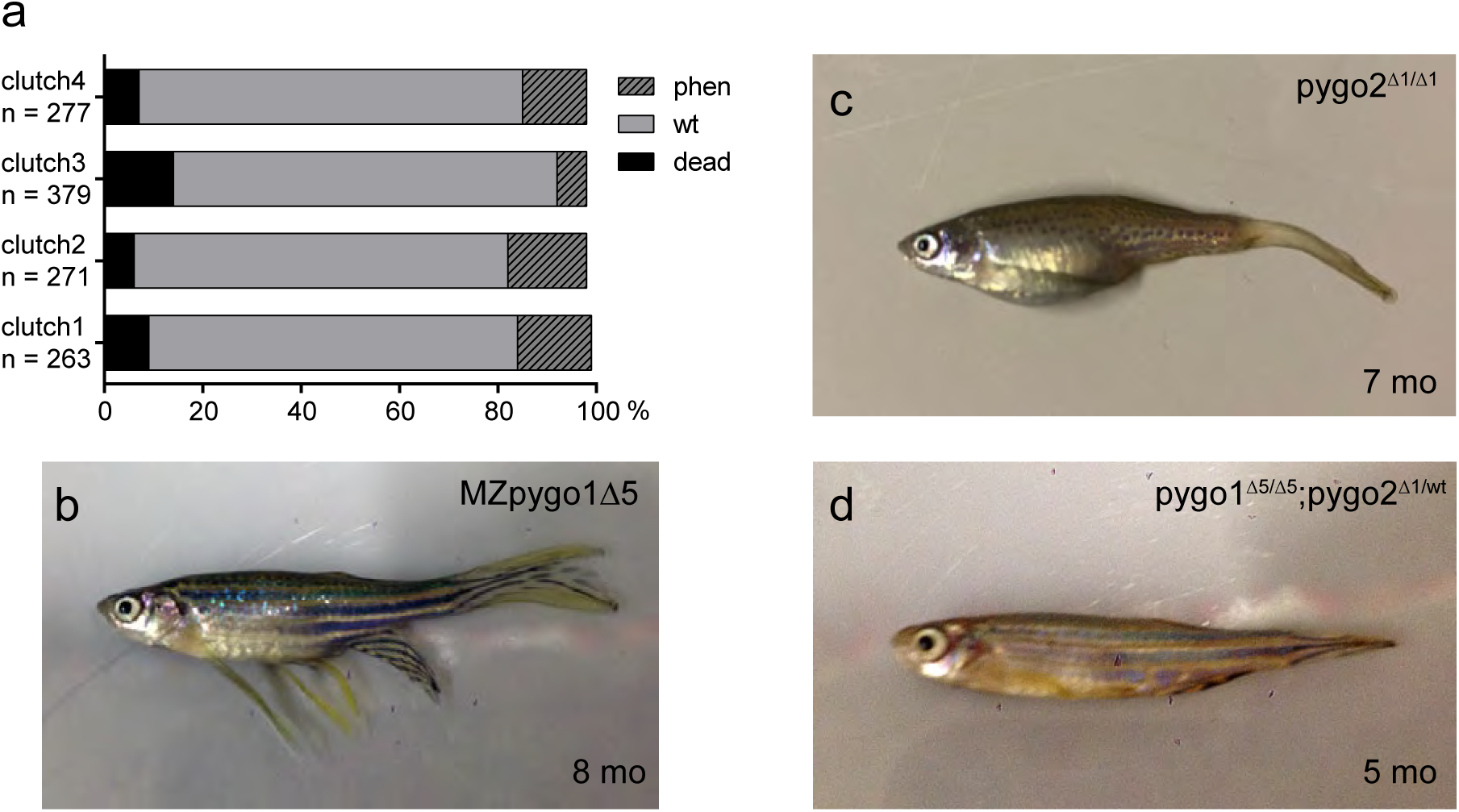
Zygotic *pygo1/2* double mutants phenocopy *bcl9* zebrafish mutants while single mutants and *pygo1* homozygous/*pygo2* heterozygous mutants are viable and fertile. (**a**) Quantification of phenotypes in four individual *pygo1^Δ5/wt^* x *pygo2^Δ1/wt^* crosses reveal defects in 7-18% of all larvae. Genotyping of phenotypic larvae revealed phenotype occurrence in homozygous *pygo2^Δ1^* mutants in combination with homozygous or heterozygous *pygo1^Δ5^* mutation. (**b-d**) MZ*pygo1^Δ5^* (**b**) and homozygous *pygo2^Δ1^* (**c**) mutants, as well as mutants carrying a homozygous *pygo1^Δ5^* combined with heterozygous *pygo2^Δ1^* alleles (**d**) are viable and fertile, lateral views, anterior to the left.

**Supplementary Figure 6:**
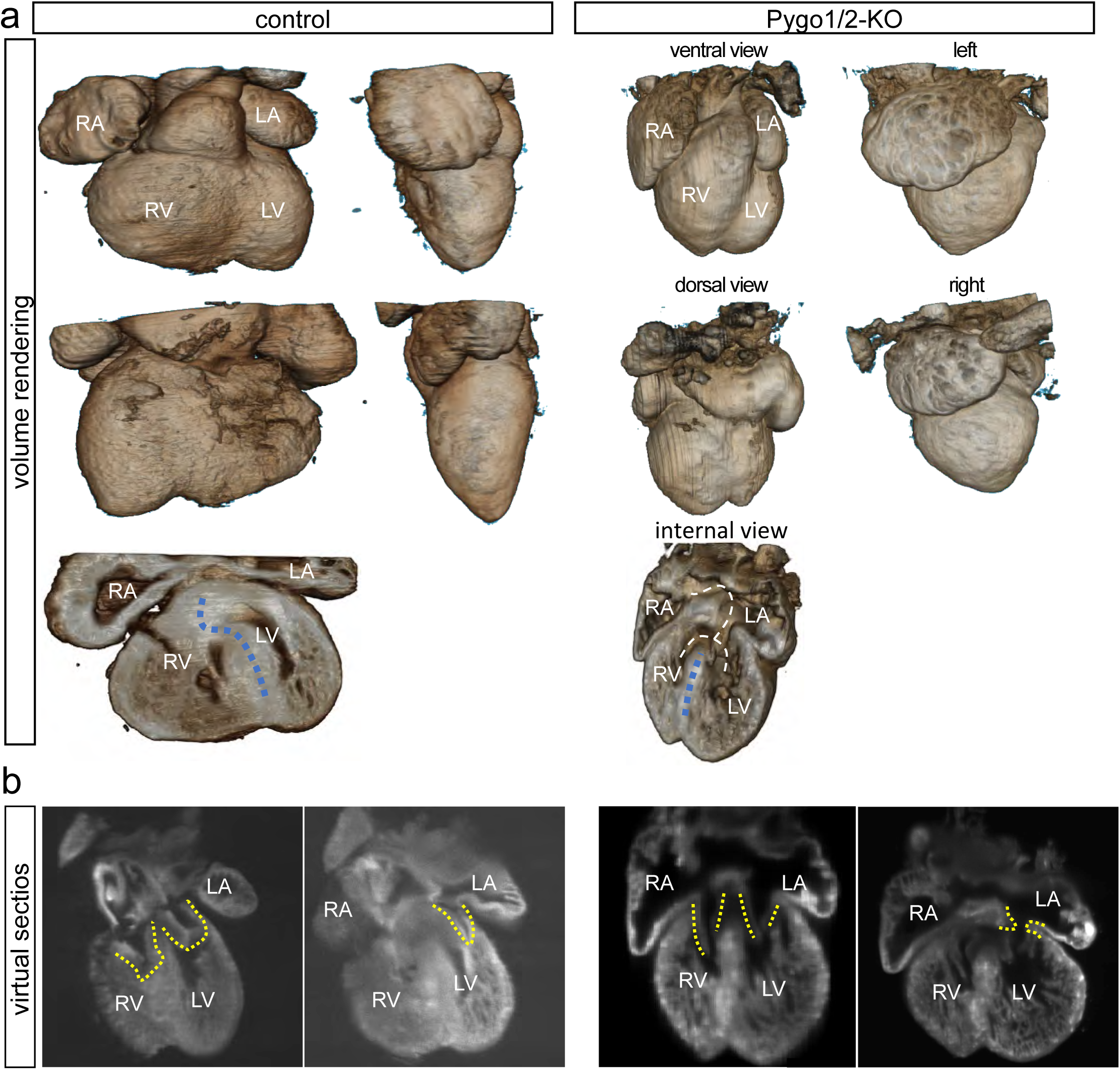
*Pygo1/2*-KO mouse embryos display severe valve and septal defects. (**a**) Volume renderings generated from SPIM images of control (left panels) and *Pygo1/2-KO* (right panels) 13.5 dpc mouse heart embryos. The images show hearts from different perspectives. Additionally, an internal view is shown on the bottom of the panel. Dashed blues lines indicate the ario-ventricular septum. Dashed white lines indicate the opening, found only in mutant hearts due to septum malformations, between the cardiac chambers. (**b**) Virtual sections generated by SPIM imaging of 13.5 dpc mouse control (left panels) or *Pygo1/2*-KO (right panels) embryonic hearts. Dashed yellow lines mark atrio-ventricular valves. Reduction in valve leaflet thickness, resulting in aberrantly communicating chambers, is evident in the mutant.

**Supplementary Figure 7:**
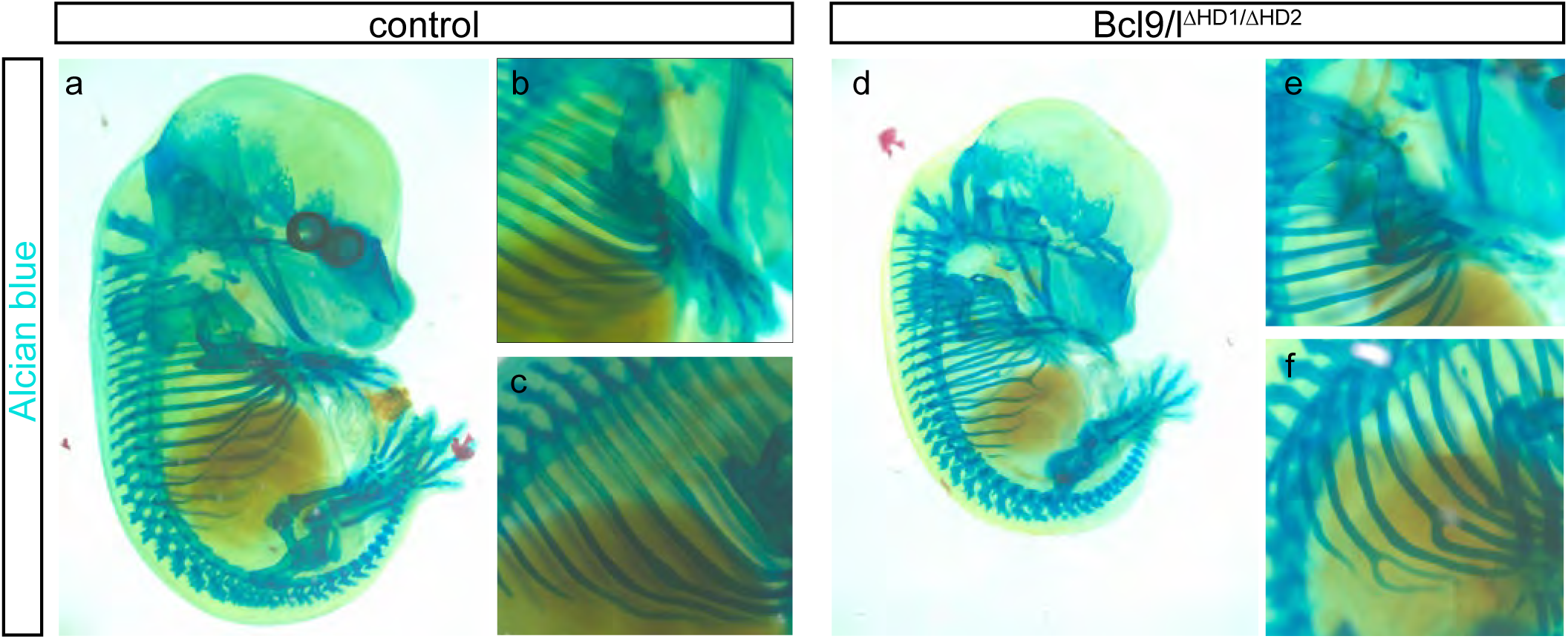
Loss of the tripartite PYGO-BCL9-β-catenin complex formation leads to cartilage defects in the mouse. (**a-f**) Alcian blue cartilage staining reveals several cartilage defects in *Bcl9/9l-Δ1/Δ2* embryos, such as loss of digit formation (compare **a,b** with **d,e**) and rib bifurcations (**c,f**).

**Supplementary Figure 8:**
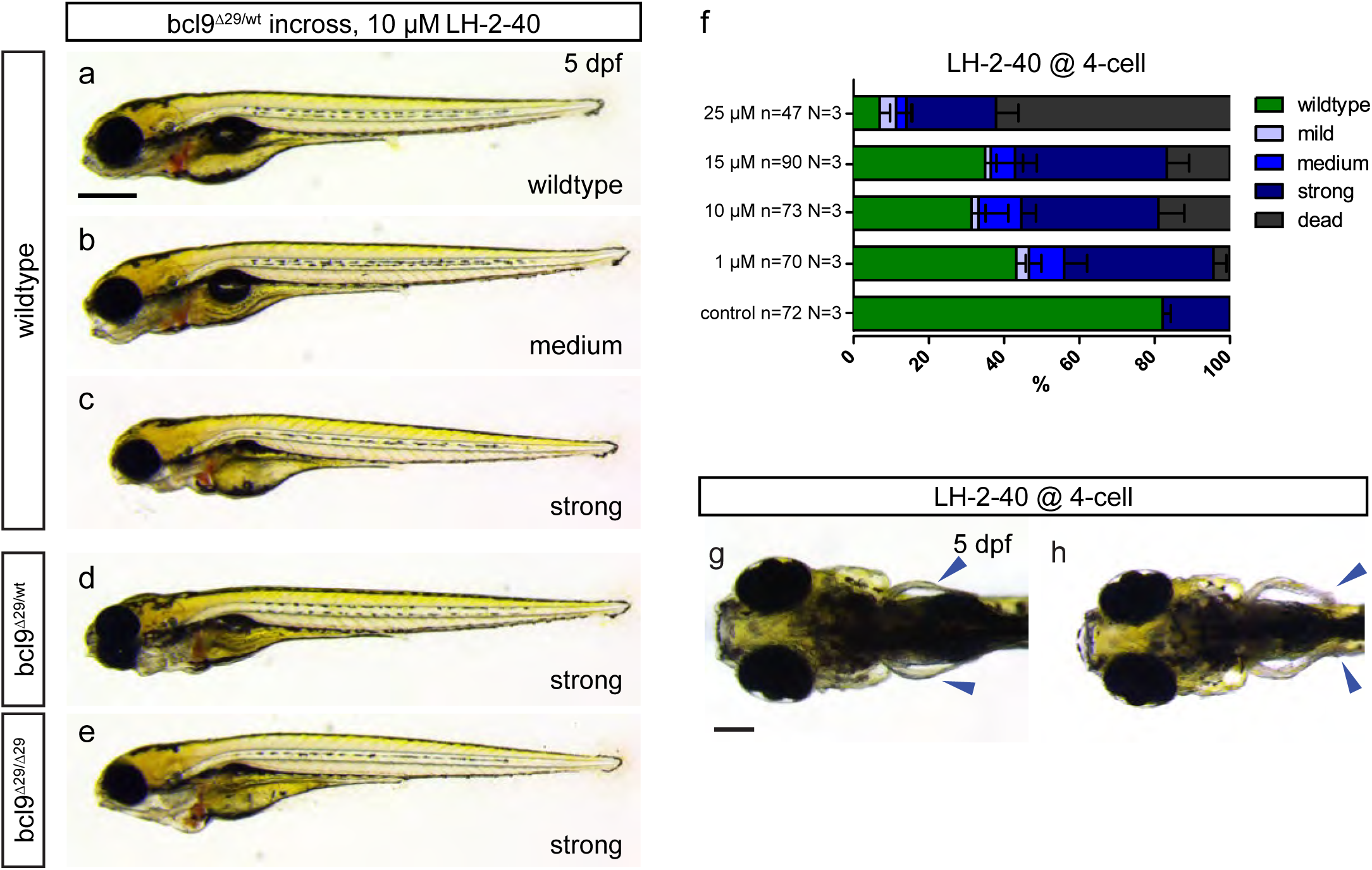
Functional inhibition of Bcl9-β-catenin-interaction in *bcl9^Δ29^* mutants and wildtypes leads to craniofacial, cardiac, and fin defects. (**a-e**) Embryos obtained from a heterozygous *bcl9^Δ29^* (*bcl9^Δ29/wt^*) incross were treated with 10 µM LH-2-40 from 4-cell stage on, lateral views, anterior to the left. Phenotype classes as described in Figure 4 were observed independent of the genotype of the larvae. (f) General phenotype penetrance is comparable to wildtype embryos treated with 10 µM LH-2-40 from 4-cell stage on (see Figure 4f). Nevertheless, the penetrance of strong phenotypes is increased in treated *bcl9^Δ29^* crosses. (**g,h**) In addition to cardiac and craniofacial defects, Bcl9-inhibited larvae (wildtypes or *bcl9^Δ29^* incrosses) are characterized by readily observable fin defects not observed in untreated *bcl9^Δ29^* mutants (arrow heads **h**, compare to control in **g**).

**Supplementary Figure 9:**
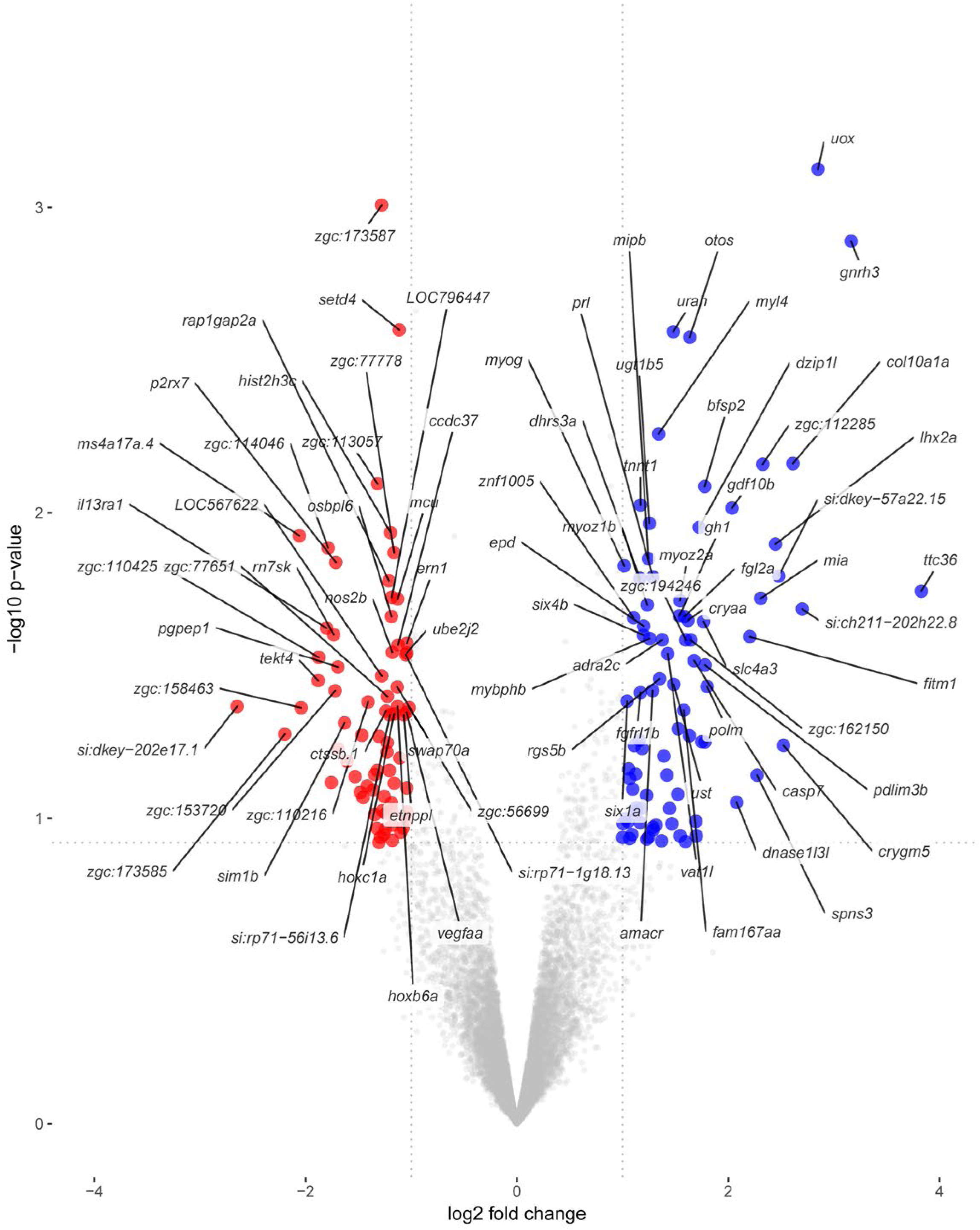
Total set of differentially expressed genes in *bcl9Δ29* mutants compared to wildtype. Volcano plot depicting the set of upregulated (74) and downregulated (83) genes in *bcl9Δ29* mutant zebrafish embryos with an absolute value of the logFC above 1, p<0.12.

**Supplementary Figure 10:**
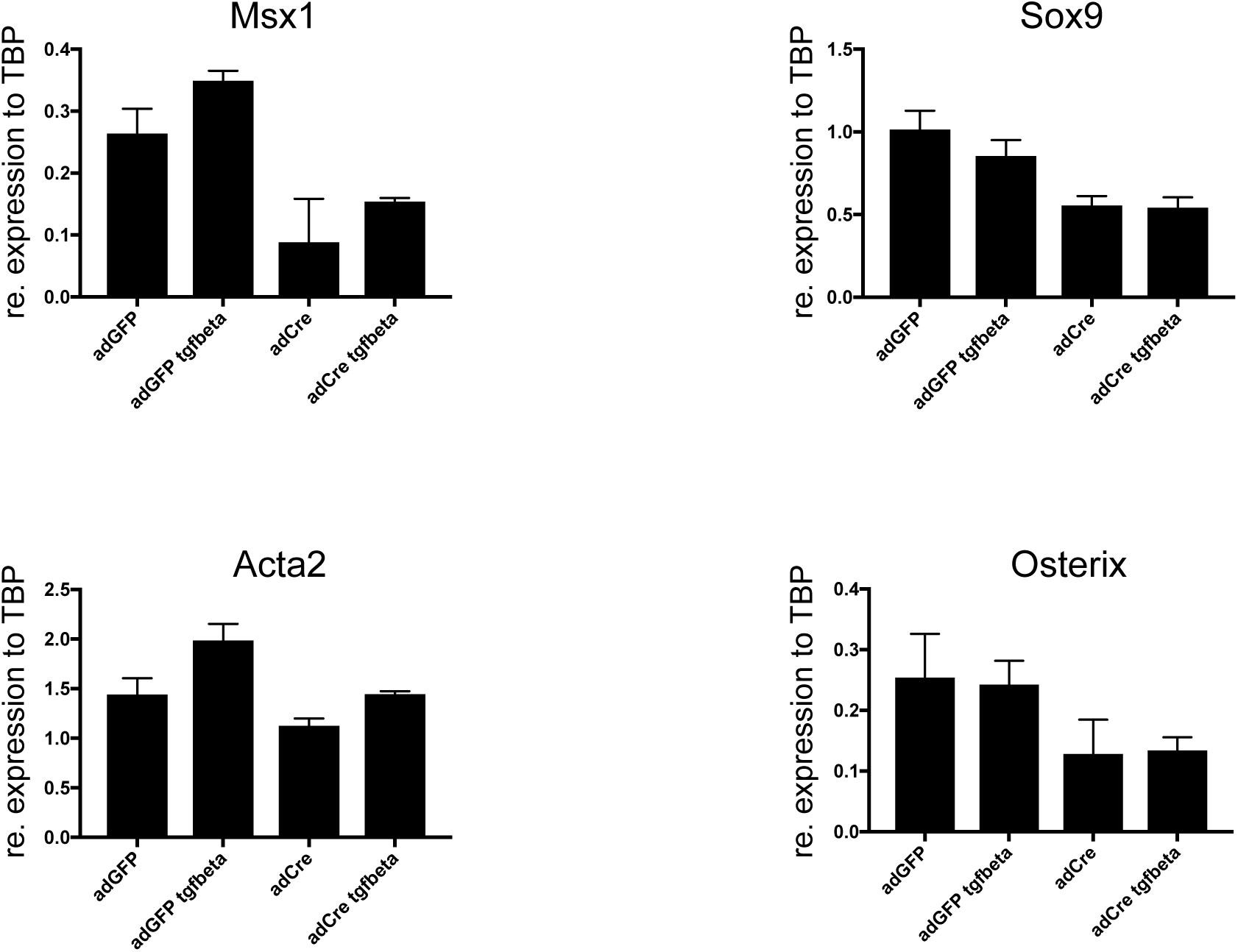
Neural crest cells deficient for Pygo1/2 exhibit reduced expression of differentiation markers. (**a-d**) Branchial arch cell explants from *Pygo1/2-flox* mice were treated with *adenoCre* viral particles and TGFβ to induce differentiation. The reduction of neural crest markers *Msx1* (**a**), *Sox9* (**b**), *SMA* (**c**) and *Osterix* (**d**) is evident upon loss of Pygo1/2.

**Supplementary Table 1.** De-regulated genes in *bcl9^Δ29^* zebrafish mutants with expression in cardiac, pharyngeal, and craniofacial precursors/derivatives as per ZFIN annotations and selected publications.

| **Gene name** | **FC** | **Zfin ID** | **Expression** |
| --- | --- | --- | --- |
| *mcu* | 0.46 | ZDB-GENE-061013-24 | heart, heart tube |
| *nos2b* | 0.44 | ZDB-GENE-080916-1 | mandibular arch skeleton |
| *zgc:56699* | 0.46 | ZDB-GENE-040426-2099 | pharyngeal arches |
| *vegfaa* | 0.49 | ZDB-GENE-990415-273 | heart, pharyngeal arch |
| *hoxc1a* | 0.45 | ZDB-GENE-000822-2 | pharyngeal arch |
| *plaub* | 0.36 | ZDB-GENE-030619-15 | heart |
| *cyp3a65* | 0.43 | ZDB-GENE-050604-1 | heart |
| *arhgap17a* | 0.31 | ZDB-GENE-050417-56 | pharyngeal arch |
| *cdh5* | 0.43 | ZDB-GENE-040816-1 | heart, heart primordium, lateral mesoderm |
| *adora2aa* | 0.46 | ZDB-GENE-080723-28 | heart |
| *rspo1* | 0.50 | ZDB-GENE-040718-44 | heart |
| *met* | 0.37 | ZDB-GENE-041014-1 | pharyngeal arch |
| *wtip* | 0.36 | ZDB-GENE-050419-261 | heart |
| *tgfbr1a* | 0.44 | ZDB-GENE-051120-75 | pharyngeal arch |
| *hoxb4a* | 0.41 | ZDB-GENE-990415-105 | pharyngeal arch |
| *vwf* | 0.40 | ZDB-GENE-070103-1 | pharyngeal arch |
| *rbck1* | 0.51 | ZDB-GENE-040704-3 | pharyngeal arch |
| *mtf1* | 0.46 | ZDB-GENE-020424-1 | heart |
| *mcoln1a* | 0.42 | ZDB-GENE-040426-2704 | heart |
| *lepr* | 0.50 | ZDB-GENE-080104-1 | heart |
| *fgfr1b* | 0.48 | ZDB-GENE-060503-14 | pharyngeal arch |
| *sema3ga* | 0.43 | ZDB-GENE-050513-3 | pharyngeal arch |
| *cldn15la* | 0.46 | ZDB-GENE-010328-12 | heart |
| *ucp3* | 0.46 | ZDB-GENE-040426-1317 | heart |
| *cldng* | 0.48 | ZDB-GENE-010328-7 | lateral mesoderm, heart |
| *erbb2* | 0.47 | ZDB-GENE-031118-121 | neural crest |
| *ltk* | 0.48 | ZDB-GENE-030616-115 | neural crest |
| *wnt1* | 0.50 | ZDB-GENE-980526-526 | neural crest |
| *matn1* | 2.68 | ZDB-GENE-050307-3 | pharyngeal arch |
| *gnrh3* | 8.96 | ZDB-GENE-030724-4 | heart |
| *pdlim3b* | 3.43 | ZDB-GENE-060130-104 | heart, heart tube, mandibular arch skeleton |
| *srfb* | 2.87 | ZDB-GENE-040426-1294 | heart rudiment, primitive heart tube |
| *fabp10a* | 3.24 | ZDB-GENE-020318-1 | heart |
| *optc* | 2.74 | ZDB-GENE-030131-1798 | pharyngeal arch |
| *myoz2a* | 2.92 | ZDB-GENE-040426-1880 | heart |

**Supplementary Table 2:** Examples of candidate genes deregulated in *bcl9^Δ29^* zebrafish mutants and with known functions in heart and craniofacial development

| Gene | in *bcl9*<sup>Δ29</sup> | Reference | Biological process |
| --- | --- | --- | --- |
| *cdh5* | down | Li et al., 2016 | Valve formation, downstream of Notch |
| *erbb2* | down | Camenisch et al. 2002 | EMT during valve formation |
| *vegfaa* | down | Lee et al. 2006 | Valve formation |
| *tgfr1a* | down | Mercado-Pimentel et al., 2007 | EMT during valve formation |
| *rbck1* | down | Wang et al., 2013 | Candidate gene for a rare CHD |
| *fgfr1b* | down | Review: Rochais 2009 | SHF/OFT development |
| *wtip* | down | Wijk et al., 2009; Powell et al. 2016 | Canonical Wnt inhibitor, epicardial development |
| *pdlim3b* | up | Pashmforoush et al., 2001 | RV cardiomyopathy, actin filament organization |
| *myoz2a* | up | Gomes et al., 2016 | Cardiomyocyte maturation, myofibril assembly |
| *myl4* | up | Review: England & Loughna 2013 | Atrial myosin |
| *sema3ga* | down | Yu et al., 2005 | Cranial neural crest cell migration |
| *wnt1* | down | Dorsky et al. 1998 | Neural crest fate induction |
| *rbck1* | down | Landgraf et al., 2010 | Craniofacial development |
| *ltk* | down | Simoes-Costa et al., 2014 | Cranial neural crest development |
| *matn1* | up | Shao et al., 2016 | Craniofacial development |

**Supplementary Table 3:** *Pygo1/2* loss-of-function leads to broad secondary gene expression changes.

| P<0.05 - GO biological process | Gene number | Expected Number | Fold Enrichment | P-value |
| --- | --- | --- | --- | --- |
| ribosome biogenesis | 71 | 18.56 | 3.83 | 5.60E-17 |
| gene expression | 325 | 207.72 | 1.56 | 1.59E-12 |
| primary metabolic process | 669 | 517.83 | 1.29 | 1.00E-11 |
| cellular nitrogen compound biosynthetic process | 289 | 181.69 | 1.59 | 1.66E-11 |
| cellular protein metabolic process | 291 | 190.65 | 1.53 | 1.81E-09 |
| ribosomal large subunit biogenesis | 29 | 5.22 | 5.55 | 3.43E-09 |
| macromolecular complex subunit organization | 170 | 95.11 | 1.79 | 3.72E-09 |
| cellular component assembly | 223 | 136.6 | 1.63 | 4.57E-09 |
| rRNA processing | 44 | 12.42 | 3.54 | 1.78E-08 |
| ribosome assembly | 26 | 4.59 | 5.67 | 3.52E-08 |
| organelle organization | 284 | 191.21 | 1.49 | 7.76E-08 |
| cytoplasmic translation | 21 | 3.03 | 6.92 | 1.11E-07 |
| RNA processing | 103 | 50.1 | 2.06 | 1.36E-07 |
| rRNA metabolic process | 45 | 13.9 | 3.24 | 1.86E-07 |
| ribonucleotide metabolic process | 56 | 20.18 | 2.78 | 2.49E-07 |
| ATP metabolic process | 39 | 11.22 | 3.48 | 5.36E-07 |
| ncRNA processing | 56 | 21.45 | 2.61 | 2.27E-06 |
| ribosomal small subunit biogenesis | 24 | 5.08 | 4.72 | 8.31E-06 |
| nucleotide metabolic process | 64 | 28.36 | 2.26 | 3.55E-05 |
| ribosomal small subunit assembly | 12 | 1.34 | 8.95 | 1.68E-04 |
| mitochondrial ATP synthesis coupled electron transport | 18 | 3.6 | 5 | 4.28E-04 |
| mitochondrion organization | 53 | 24.13 | 2.2 | 1.69E-03 |
| regulation of programmed cell death ( | 154 | 102.52 | 1.5 | 4.35E-03 |
| cellular localization | 187 | 132.16 | 1.41 | 1.17E-02 |
| regulation of cell death | 162 | 111.91 | 1.45 | 1.82E-02 |
| regulation of gene expression | 341 | 270.03 | 1.26 | 2.03E-02 |
| regulation of apoptotic process | 149 | 101.32 | 1.47 | 2.12E-02 |
| muscle system process | 37 | 16.09 | 2.3 | 4.12E-02 |

